# Whole-Brain Models of Advanced Concentrative Absorption Meditation: Approaching Critical Dynamics through Jhāna

**DOI:** 10.1101/2025.09.25.678574

**Authors:** Jakub Vohryzek, Edmundo Lopez-Sola, Winson F. Z. Yang, Yonatan Sanz Perl, Ruby M. Potash, Ruben E. Laukkonen, Terje Sparby, Morten L. Kringelbach, Giulio RufJini, Gustavo Deco, Matthew D. Sacchet

## Abstract

Advanced meditation offers a powerful lens for investigating consciousness and for understanding how sustained training may contribute to human flourishing. In the spirit of neurophenomenology, we combine first-person reports with model-free empirical analyses and formal whole-brain modeling to investigate the mechanisms underlying advanced meditative states and minimal phenomenal experience (MPE). Specifically, we focus on jhāna meditation, a type of advanced concentrative absorption meditation (ACAM-J). Advanced practitioners accessed the eight ACAM-J states during ultra-high-field 7T functional magnetic resonance imaging. For each state, we first characterize empirical functional connectivity and then build a mechanistic whole-brain model that reproduces brain activity by modeling the dynamical regimes of different brain networks. We found that the later ACAM-J states, taken here as candidates for MPE, show increased large-scale functional integration and a shift of functional network dynamics toward near-critical working points. The default mode network (DMN) exhibits the largest shift, from a distant noise-driven regime during the control condition to near-critical dynamics during ACAM-J. We also observed that the trajectory of ACAM-J states is non-linear, with prominent reconfigurations at key meditative milestones. Our results suggest that MPE, as instantiated in later ACAM-J states, corresponds to a globally susceptible state where near-critical dynamics dominate. We interpret this near-critical regime as a form of “openness”, in which constrained and differentiated patterns of brain activity give way to greater flexibility. In particular, increased DMN susceptibility is correlated with broader attention and reduced narrative thought, consistent with a more 8lexible mode of self-related processing. In this context, advanced meditation provides a powerful model for studying how sustained contemplative practice can profoundly shape brain dynamics and provide a window into core aspects of consciousness.

## Introduction

Advanced meditation provides a unique opportunity to investigate consciousness (Lieberman et al., 2025; Sacchet et al., 2026), including its relationship to well-being, meaning-making, and human flourishing (Kringelbach et al. 2024). Unlike altered states of consciousness induced by external perturbations such as pharmacological drugs, advanced meditative states are volitional, stable, and reproducible by experienced practitioners (Ganesan et al. 2024). Their systematic progression enables mechanistic investigation, wherein neural activity during meditation can be interpreted in relation to precisely characterized phenomenology, as recommended by Varela (1996) under the paradigm of neurophenomenology.

Within this context, *jhāna* practice, a type of advanced concentration absorption meditation in Buddhist contemplative practices, comprises a series of eight meditative absorption states with relatively well-defined phenomenological qualities. Contemporary practice manuals provide detailed phenomenological descriptions (Brahm et al. 2006; Brasington 2015; Burbea 2014; Ingram 2018), and recent theoretical work has proposed a more systematic classification of the jhānas as a type of advanced concentrative absorption meditation (ACAM-J; Sparby et al. 2024), as part of a “third wave” of meditation research that focuses specifically on advanced meditative states (Sacchet et al., 2024).

ACAM-J unfolds progressively, beginning with effortless, stable focus that synthesizes each moment of experience, creating continuity and absorption (J1). This experience leads to the unfolding of a series of experiences of unification that proceed from intense bliss and happiness, a kind of peak experience (J2), to contentment (J3) and increasing mindfulness, peace, and equanimity (J4). Beyond these four (*form*) ACAM-J, further potential refinements of consciousness may ensue as other contents of experience (e.g., sensory input, thoughts, affect) are muted or disappear completely. Minimal and yet expansive structural features of experience may come into focus, including minimal expansive spatiality (J5) and minimal expansive consciousness (J6). The most subtle stages, absence perception (J7) and liminality perception (J8), involve extremely sparse modes of awareness approaching liminal or near-absent phenomenology.

Particularly relevant to this study is the later ACAM-J as they are perceptually minimal (Sparby et al. 2024). In these states, sensory impressions fade, subject–object structure weakens, and experiential content becomes increasingly attenuated (Burbea 2014; Brasington 2015; Ingram 2018). Notably, these phenomenological features closely converge with accounts of minimal phenomenal experience (MPE) collected in Metzinger’s recent compilation of the general phenomena (Metzinger 2024) which aims to identify structural features sufficient for conscious experience (Metzinger 2020). We draw on this framework to test the later ACAM-J as potential empirical approximations of MPE. Because progression through ACAM-J is structured and reproducible, it provides investigation of the neural mechanisms underlying the gradual reductions in experiential structure.

In line with neurophenomenological approaches (Lutz et al. 2025; Thompson et al. 2005), recent computational accounts suggest that ACAM-J reflects a reduction of hierarchical modeling, (Laukkonen et al. 2023; Lopez-Sola et al. 2025; Mago et al. 2024; Prest et al. 2024), described phenomenologically as a process of “letting go,” and formally modeled as a progressive reduction of prior precision or as the dissolution of attractor states along the modeling hierarchy, potentially accompanied by increased entropy (Mago et al. 2025). However, recent studies on ACAM-J have yet to investigate this using computational models that reproduce the underlying brain mechanisms (e.g., Yang et al. 2024; Ganesan et al. 2024, Demir et al. 2025; Potash et al. 2025, Yang et al. 2025b).

Whole-brain models provide such a framework by combining formal descriptions of local neural dynamics with anatomically constrained structural connectivity (Deco et al. 2013) that enables reproduction of a wide range of phenomena, including resting-state functional connectivity, metastability, and signal complexity (Breakspear 2017; Deco et al. 2017). Whole-brain models have already been applied to a variety of states of consciousness, including sleep, anesthesia, disorders of consciousness, and pharmacologically-induced states (Deco et al. 2018a, 2018b; Gendra et al. 2024; Herzog et al. 2023; López González et al. 2021; Luppi et al. 2022; Mindlin et al. 2024; Sanz Perl et al. 2021; Vohryzek et al. 2024b) where changes in entropy and functional hierarchy can be characterized.

To date, only two studies have so far applied large-scale models to meditative states (Dagnino et al. 2024; De Filippi et al. 2022). Both examined a hybrid concentration-insight practice (vipassanā *anapanasati*), which combines concentration and insight but follows a different phenomenological pathway than the later ACAM-J states. Here, we extend large-scale modeling to later ACAM-J to characterize the dynamical regimes associated with perceptually minimal states.

A closely related organizing concept within this modeling framework is *criticality*. Criticality refers to a dynamical regime poised at the transition between ordered and disordered activity, where system dynamics are neither rigid nor random, but maximally responsive to internal and external influences. Empirical and modeling studies suggest that the brain operates near this critical point during wakefulness, whereas departures are observed during states such as deep sleep, anesthesia, or certain pathological conditions (Cocchi et al. 2017). At this regime, small perturbations can propagate across the system, amplifying their effects and increasing complexity, while still maintaining enough stability to support a coherent function (Figure 1). Whole-brain models frequently achieve optimal fittings near bifurcation points, balancing between order and instability that supports both consistency and adaptability (Deco et al. 2012; Sanz Perl et al. 2021, 2022).

**Figure 1:**
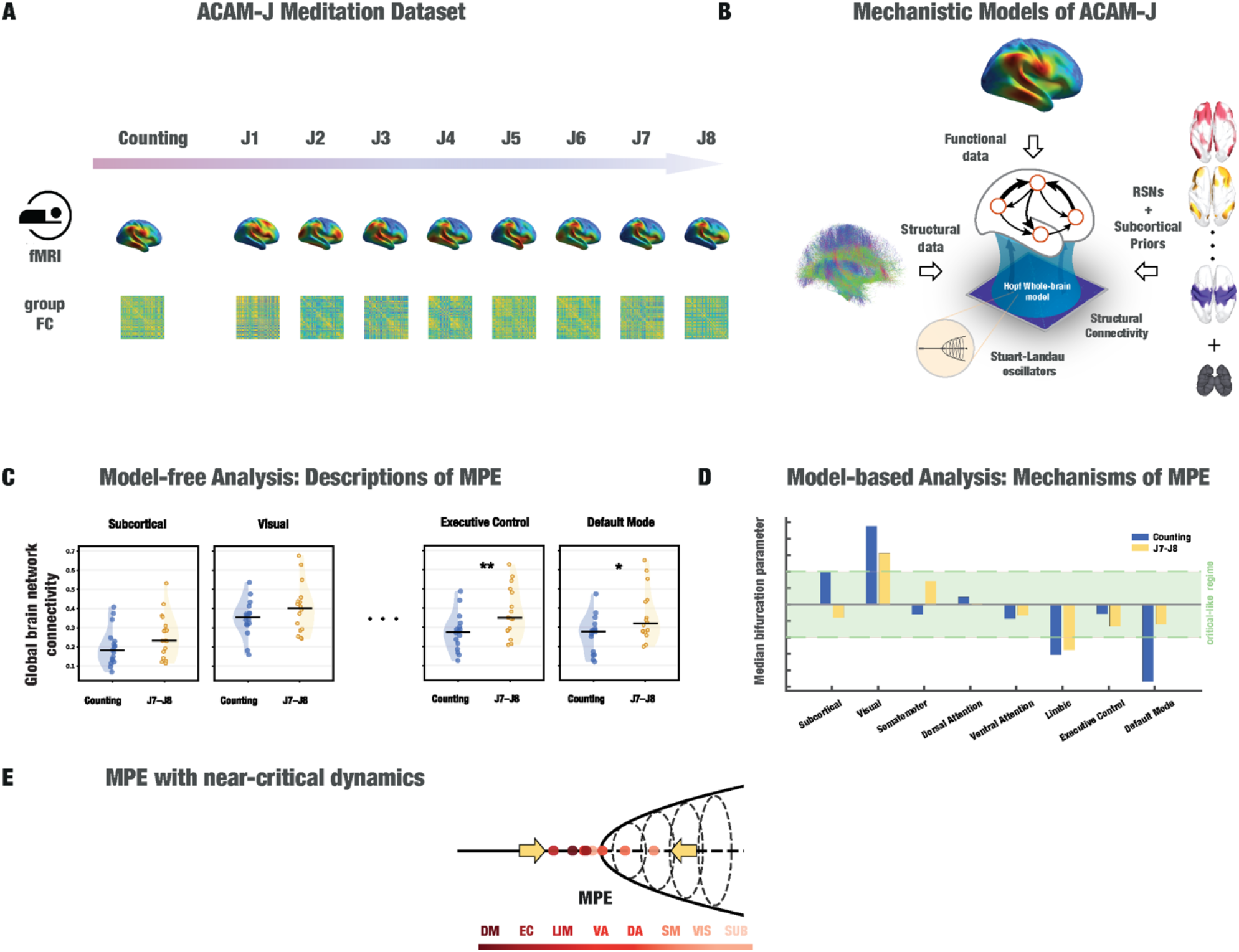
Investigation of mechanisms driving ACAM-J. **A:** Dataset description: we used fMRI data of 20 subjects undergoing ACAM-J. We computed a group functional connectivity (FC) to fit whole-brain models for each of the 8 ACAM-J states and the control counting condition. **B:** We created whole-brain models to fit the group FC. Whole-brain models, based on coupled local dynamics reflecting Stuart-Landau oscillators and constrained by a normative structural connectome, were fitted to the group FC of each condition. We used the resting-state networks defined in Yeo et al. (2011) as spatial priors for a genetic algorithm that fitted the bifurcation parameter of each local oscillator. **C:** Model-free Analysis: We quantified the MPE state in terms of global brain connectivity across major networks by comparing Counting with late Jhana practice, summarized as mean ACAM-J7-J8. **D:** Model-based Analysis: We found that for most RSNs the bifurcation parameter converged towards near-critical dynamics in MPE (here represented by ACAM-J7-J8). **E:** We discuss the hypothesis that MPE is characterized by a collapse of the brain’s dynamical regimes towards near-critical dynamics.

Recent theoretical work has proposed that advanced meditation and MPE may involve shifts to critical dynamics and increased complexity (Mago et al. 2024; Sandved-Smith 2024). This proposal is consistent with descriptions of meditation as avoiding both excessive rigidity (e.g., fixation, habitual reactivity) and instability (e.g., agitation, loss of control). This is further supported by emerging empirical studies in deep meditative states (Demir et al. 2025; van Lutterveld et al. 2025). Computational models are particularly well-suited for testing this hypothesis because they make the relevant control parameters explicit by tuning local or global parameters that govern the stability of regional dynamics (Cocchi et al. 2017).

In this work, “near-critical” refers to proximity to a supercritical Hopf bifurcation, where an oscillatory mode becomes weakly damped, yielding increased responsiveness without implying thermodynamic phase transitions or scale-free statistical behavior (Bose et al., 2019). Computational modeling thus provides a powerful way to study criticality as a dynamical property, which is a key framework for understanding how different conscious states, including deep meditative absorption, reorganize large-scale brain dynamics. Framing ACAM-J within this dynamical context allows investigation of how large-scale brain activity reorganizes as experiential structure approaches minimality.

In the current study, we first quantified model-free global brain network connectivity to characterize large-scale functional integration directly from the empirical data. Then, we created generative whole-brain models of large-scale functional organization across ACAM-J and its relevant perceptually minimal states. Whole-brain models were fitted to fMRI data from 20 experienced practitioners across ACAM-J and a control condition, with dynamics parameterized by canonical resting-state networks (RSNs). Crucially, model parameters were examined in relation to graded phenomenological reports, enabling direct linkage between dynamical regime shifts and phenomenology. This framework allows assessment of how differential RSN involvement supports distinct ACAM-J and whether progression through perceptually minimal states is accompanied by systematic shifts to near-critical dynamics.

## Methods

### Participants

The present dataset included 20 advanced ACAM-J practitioners and is fully described in Yang et al. (2025b). Group-level brain states were estimated using all available recordings per state; consequently, sample sizes varied across states, with 19 subjects contributing to J3, J5, and J6, 18 to J7, and 17 to J8. All participants provided informed consent, and ethical approval was obtained from the relevant institutional review board. Similar to the original study, participants were screened for neuropsychiatric and cognitive health to ensure suitability for high-field fMRI research.

### Meditation Protocol

Each participant performed a standardized sequence of ACAM-J inside the MRI scanner. The sequence involved entering access concentration (Brasington 2015), progressing systematically through the first to eighth ACAM-J (J1–J8), and concluding with a post-jhānic “afterglow” state. As in Yang et al. (2025b), transitions into each ACAM-J state were indicated using button presses. Neuroimaging acquisition parameters and preprocessing details are provided in the Supplementary Information.

### Control Conditions

To provide non-meditative comparison conditions, participants completed two tasks designed to prevent inadvertent entry into meditative absorption: *Memory recall* (mentally recounting personal events from the previous two weeks) and *Mental counting* (silently counting backward in decrements of five from a large starting number, e.g., 10,000). Each control task lasted approximately 8 minutes. Runs were scheduled both before and during meditation sessions, as in the original design (Yang et al. 2025b).

In the present study, we focus on the counting condition as the primary control. Empirically, this condition yielded results that more closely matched prior whole-brain modeling findings in wakeful resting-state than the memory condition, including the presence of subcritical dynamics in higher-order networks (for example, default mode and executive control) and supercritical dynamics in lower-order sensory systems (for example, visual and somatomotor cortex) (Ipiñ a et al. 2020, Perl et al. 2023). To further characterize the baseline conditions, we directly compared the dynamical regimes obtained under the counting and memory tasks (see Figure S6). These control conditions are not assumed to be equivalent but instead probe distinct cognitive regimes: counting approximates a task-constrained baseline consistent with canonical resting-state whole-brain dynamics, whereas memory recall engages internally generated, autobiographical processing. Given that our hypothesis concerns deviations from a baseline regime aligned with resting-state dynamics, the counting condition was selected as the primary reference.

### Global brain network connectivity

To quantify large-scale functional integration, we computed global brain network connectivity for each resting-state network (RSN) and the subcortical group using empirical fMRI BOLD data. For each subject and condition, functional connectivity matrices were obtained by computing pairwise Pearson correlations (Fisher z-transformed) between all brain regions. These region x region matrices were then averaged within and between predefined RSNs and the subcortical group to obtain an RSN+1 x RSN +1 connectivity matrix. For each RSN and the subcortical group, global connectivity was defined as the average across its corresponding row (i.e., its connectivity to all networks, including itself), yielding a single value per network, subject, and condition.. Group-level distributions were visualized using violin plots. Statistical comparisons between conditions were performed using both parametric (paired t-tests) and non-parametric (Wilcoxon signed-rank tests) approaches, with p-values corrected for multiple comparisons using false discovery rate (FDR).

### Whole-Brain Model

We implemented a Hopf bifurcation model to describe local neural mass dynamics. Each brain region *j* was modeled as:

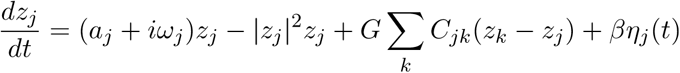

where *z*_*j*_ = *x*_*j*_ +*iy*_*j*_ represents regional activity, *a*_*j*_ is the bifurcation parameter, *ω*_*j*_ the intrinsic frequency (estimated from fMRI peak frequency), *C*_*jk*_ the structural connectivity weight, *G* a global coupling parameter, and *βη*_*j*_ Gaussian noise. The bifurcation parameter represents different dynamic regimes of the Hopf oscillator — *a*_*j*_ *<* 0: stable fixed-point (noise-driven), *a*_*j*_ *>* 0: self-sustained oscillations, *a*_*j*_ ≈ 0: complex, metastable dynamics. Details of the structural connectivity construction are provided in the Supplementary Information.

In the present study, proximity to the Hopf bifurcation (near-critical regime) was operationally defined as -0.02<a<0.02. Values within this narrow band correspond to weakly damped or weakly unstable dynamics with long relaxation times and increased susceptibility to perturbations. Previous whole-brain modelling studies have shown that optimal fits to empirical resting-state activity typically cluster within this range around the bifurcation (Deco et al. 2017; Sanz Perl et al. 2021).

### Optimization

In order to mechanistically test the influence of different resting-state networks in ACAM-J, regional bifurcation parameters were grouped using spatial priors to reflect seven canonical resting-state networks (Yeo et al. 2011), Visual (VIS), Somatomotor (SM), Dorsal Attention (DA), Ventral Attention (VA), Limbic (LIM), Executive Control (EC), and Default Mode (DM), and a group of subcortical regions, which we will loosely name the Subcortical group (SUB).

First, we fixed the global coupling to G = 0.5, as optimized in previous literature (Ipiña et al. 2020). Spatial heterogeneity in local dynamics was introduced by grouping brain regions according to anatomically motivated priors (in this case, the mapping of the eight networks into the Schaefer-200 (N=200) + Tian Scale 2 (N=34) parcellation), with each group contributing an independent coefficient to the bifurcation parameter of its nodes. These coefficients were estimated using a genetic algorithm that iteratively improved candidate solutions based on their fit to the empirical data. Implementation details of the genetic algorithm are provided in the Supplementary Information. The empirical observables were derived from averaging participants per condition (across fourteen subjects), producing group-level FC matrices (calculated in terms of pairwise Pearson correlation) for each ACAM-J. Fitness was quantified as one minus the structural similarity index (SSIM) between empirical and simulated connectivity matrices.

We also ran a homogeneous optimization, where we optimized a global bifurcation parameter. Unlike the heterogeneous case, where we parametrized the bifurcation parameter of each RSN, here we optimized only one bifurcation parameter.

### Statistical Analysis

To contrast model outputs across conditions, we generated forty independent realizations and derived the corresponding distributions of values. Differences between conditions were quantified using Cohen’s d, which captures the magnitude of effect size. As in previous work (Perl et al. 2023), we do not report p-values or other significance metrics dependent on sample size, since the apparent statistical power could be inflated arbitrarily by increasing the number of model realizations. For clarity, we follow conventional interpretations of Cohen’s d in which values lower than 0.2 indicate small effects, values around 0.5 reflect medium effects, and values of 0.8 or above correspond to large effects (Cohen 2013).

### Phenomenological reports

A structured first-person phenomenological assessment was conducted following completion of the full ACAM-J sequence (ACAM-J1 to ACAM-J8). Participants provided ratings on a 1 to 10 Likert scale. For each control condition and ACAM-J state, participants reported stability of attention and width of attention. For each ACAM-J state, participants rated the overall quality of that ACAM-J, and the specific ACAM-J factor associated with that state, namely bliss or joy (ACAM-J1–2), contentment (ACAM-J3), equanimity (ACAM-J4), and formlessness (ACAM-J5–8). In addition, the presence of experiential contents, including sights, sounds, physical sensations, and narrative thought stream, was rated separately for the aggregate early (ACAM-J1–4) and late (ACAM-J5–8) states. Higher values indicated greater intensity or prominence of the corresponding feature.

To test whether the dynamical regime of the DMN tracked inter-individual differences in phenomenological features, we adopted a two-level summary-statistics approach. For each participant and each phenomenological feature, we computed Pearson’s correlation coefficient between the condition-wise DMN bifurcation parameter distance to the bifurcation (|a|) and that participant’s phenomenological ratings. Because some phenomenological dimensions were rated at the block level (early and later ACAM-J aggregates), DMN bifurcation values were collapsed accordingly to three levels, counting, mean(J1–J4), and mean(J5–J8), to ensure matched resolution and avoid pseudo-replication. For the ACAM-J factor rating, only the formlessness factor (J5–J8) was analyzed, as it was the only factor rated across enough conditions per participant (4) to support a within-subject correlation; the remaining factors spanned too few conditions (bliss/joy: 2; contentment: 1; equanimity: 1). This yielded one correlation coefficient per subject per feature. At the second level, the distribution of per-subject correlation coefficients was tested against zero using both a one-sample t-test and a Wilcoxon signed-rank test. *P*-values were corrected for multiple comparisons across the phenomenological features using the Benjamini–Hochberg false discovery rate (FDR) procedure. Effects were considered significant at FDR-corrected *q* < 0.05.

As a complementary analysis, we pooled all data points across participants and conditions to assess group-level associations between DMN bifurcation distance and phenomenological ratings. For each feature, we computed the Spearman rank correlation and fitted a linear mixed-effects model (rating ∼ distance to bifurcation + (1 | participant)) using restricted maximum likelihood (REML), which accounts for the non-independence of repeated observations within participants. P-values for the fixed effect of bifurcation were corrected across features using the Benjamini–Hochberg FDR procedure.

## Results

To investigate neural mechanisms underlying the progression through ACAM-J, we analyzed fMRI BOLD data from 20 advanced practitioners scanned during the counting control condition and across the eight successive ACAM-J states (**Figure 1A**). For each condition, we first computed model-free global brain network connectivity within each resting-state network and the subcortical group, allowing us to characterize empirical changes across the progression and between control and late ACAM-J states. We then computed the group-level functional connectivity and then fitted a whole-brain model of coupled Stuart–Landau oscillators (a whole-brain Hopf model), constrained by normative structural connectivity and informed by priors from the seven canonical resting-state networks and a group of subcortical regions (**Figure 1B**). Using a genetic algorithm, we optimized the bifurcation parameters of the model to reproduce the empirical functional connectivity patterns for each state, yielding network-level dynamical regimes across counting and ACAM-J states. This parametrization allowed us to characterize the dynamical regime of each resting-state network in relation to the bifurcation point, comparing the late ACAM-J with the control condition (**Figure 1D**). By comparing these network-level bifurcation parameters across the progression of ACAM-J states, we identified characteristic trajectories of RSN dynamical shifts (**Figure 1E**). Finally, we discuss the implications of these converging network dynamics for understanding how the brain approaches critical dynamics as practitioners progress through ACAM-J states (**Figure 1F**).

### Changes in dynamical regimes in MPE

By comparing the control condition with the latest ACAM-J states (J7–J8) as states representative of MPE, we identified systematic large-scale reconfigurations across RSNs and the subcortical group.

At the empirical level (Model-free Analysis), global brain network connectivity revealed an overall increase in functional integration in MPE states (**Figure 2A**). Most resting-state networks showed higher global brain network connectivity in ACAM-J7-J8 compared to the counting control condition. In particular, the Subcortical group, and the Ventral Attention, Executive Control, and Default Mode networks exhibited significant increases (Subcortical: t(16) = 2.75, p_FDR = 0.033; Ventral Attention: t(15) = 2.71, p_FDR = 0.033; Executive Control: t(15) = 4.00, p_FDR = 0.009; Default Mode: t(15) = 3.58, p_FDR = 0.011) consistent across both parametric and non-parametric analyses. Other networks followed the same increasing trend but did not reach statistical significance. Acknowledging that the subcortical group consists of functional variable regions, we conducted a follow-up subcortical-region analysis that showed that the subcortical group effect was not confined to a single subcortical subdivision. When each bilateral subcortical region was summarized by its equal-weighted connectivity to the seven cortical RSNs, most regions showed positive J7-J8 shifts, with the strongest raw increase in posterior globus pallidus, although no individual subcortical region survived FDR correction (**Figure S8**). Overall, these results indicate a widespread, though heterogeneous, increase in large-scale integration during MPE.

**Figure 2:**
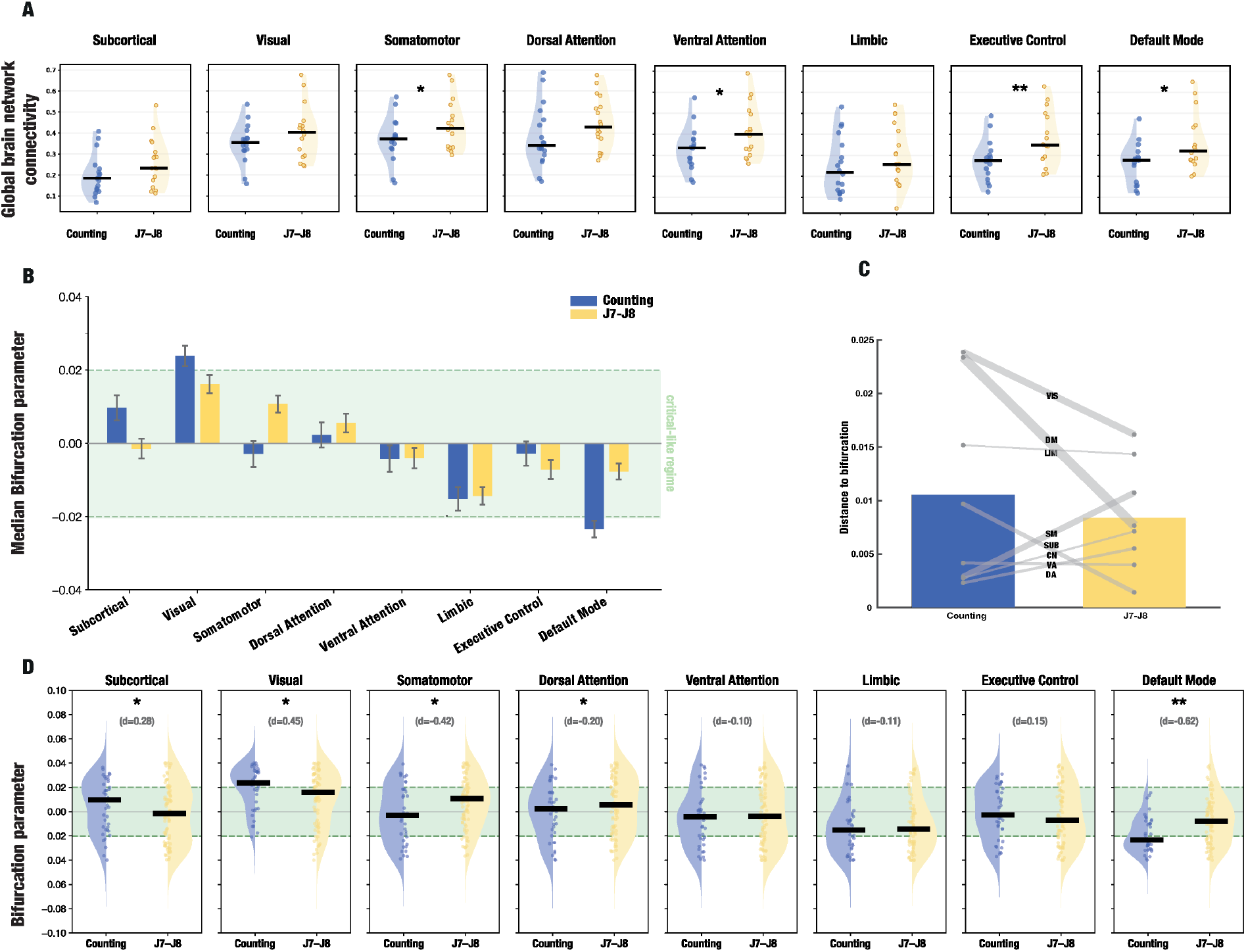
Global network connectivity increases and network dynamical regimes shift to near-critical dynamics in MPE. **A:** Distribution of global network connectivity values per RSN. Individual dots represent subject values, solid lines indicate median values across subjects, and an asterisk indicates significance in FDR-corrected t-test and signed-rank tests. Solid lines indicate median values and **\*** and **\*\*** indicate FDR-corrected statistical significance p<0.05 and p<0.01 respectively. **B**: Median bifurcation parameter values per RSN in the control and latest ACAM-J (J7-J8) conditions. The green shaded area marks the near-critical region (|*a*| *<* 0.02). **C**: Median distance of RSNs to the bifurcation point (*a* = 0). Dots indicate median distances per RSN, and line width represents the effect size according to Cohen’s d. **D**: Individual dots represent bifurcation parameters per RSN for forty runs of the genetic algorithm in each condition. Solid lines indicate median values, **\*** and **\*\*** indicate weak (0.2 *<* |*d*| *<* 0.5) and moderate (0.5 *<* |*d*| *<* 0.8) effect sizes, respectively, according to Cohen’s d.

We next asked how these large-scale changes in functional integration relate to underlying dynamical regimes using whole-brain modeling (Model-based Analysis). In the control condition, RSNs showed clear differences in dynamical regimes and in the bifurcation parameter distance to the bifurcation point (**Figure 2 and Figure S6**). In particular, the DM was relatively far from the bifurcation point in both control conditions, situated in the noise-driven regime.

When comparing the ACAM-J7/J8 to the control condition, many RSNs exhibited important changes (weak to moderate effect sizes), and these changes were generally in the direction of approaching the bifurcation point (**Figure 2C**). Importantly, in the ACAM-J7/J8 states, all networks exhibited near-critical dynamics. The Default Mode network showed the strongest shift to the bifurcation point, with moderate effect sizes (*d >* 0.5). The Subcortical group and the Visual network also moved to the bifurcation point, but with weaker effect sizes (*d* = 0.28 and *d* = 0.45, respectively). The Somatomotor network shifted from noise-driven to oscillatory dynamics with a weak effect size (*d* = −0.42) but remained close to the bifurcation point across both conditions. The Dorsal Attention, Ventral Attention, Limbic, and Control networks exhibited negligible effect sizes, all remaining close to the bifurcation point (**Figure 2D**). Importantly, this trend was replicated when using the later ACAM-J as candidate MPE states, confirming the robustness of these findings (**Figure S1**). Notably, the direction of the effect in the Default Mode network was preserved when using the memory condition, whereas other networks showed less consistent effects across control conditions (**Figure S6**). In summary, the shift to critical dynamics was evident across several networks, but its magnitude varied, reflecting heterogeneous trajectories.

Among all RSNs, the Default Mode network underwent the strongest shift between conditions. In the control condition, the DM was relatively far from the bifurcation point with a negative value of the bifurcation parameter, indicating that its dynamics were deep in the noise-driven regime (median *a* = −0.023). However, in ACAM-J7/J8 the DM was markedly closer to the bifurcation point, showing the largest effect size across all networks (*d* = -0.62). We will comment on a potential interpretation of such shifts in the discussion.

### The progression through ACAM-J

To investigate the brain mechanisms involved in the progression through ACAM-J, we examined the changes in RSNs bifurcation parameters across counting, early ACAM-J (J1–J4), and late ACAM-J (J5–J8) states. Results reveal a non-linear approximation towards near-critical dynamics, characterized by stronger reconfigurations in the early ACAM-J and a smoother convergence in later states (**Figure 3**).

**Figure 3:**
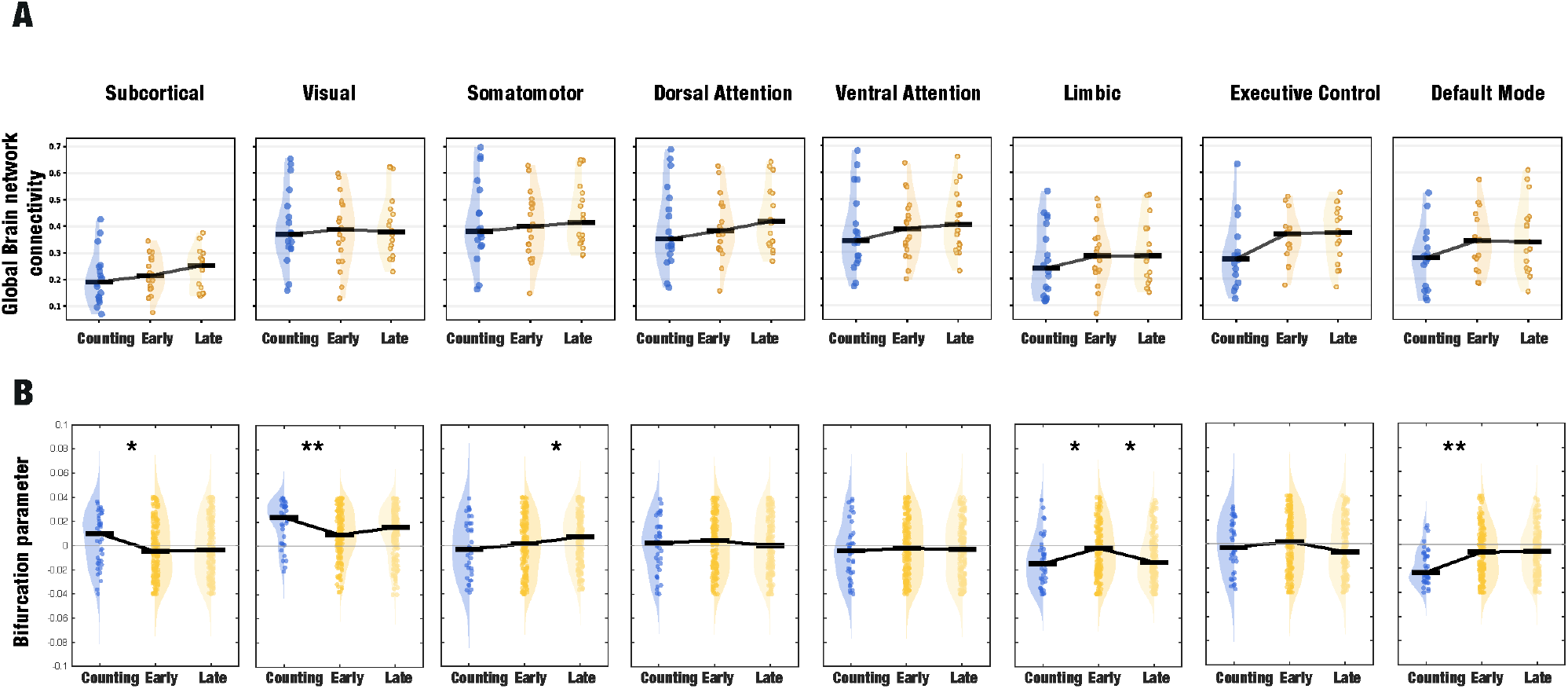
Progression to MPE through early and late ACAM-J states. **A:** Global brain network connectivity values per RSN in the counting, early ACAM-J (J1-J4) and late ACAM-J (J5-J8) conditions. Individual dots represent subject values, solid lines indicate median values across subjects, and an asterisk indicates significance in FDR-corrected t-test and signed-rank tests. **B:** Individual dots represent the parameters per RSN for forty runs of the genetic algorithm in each condition (counting, early and late ACAM-J states). Solid lines indicate median values, **\*** and **\*\*** indicate weak (0.2 *<* |*d*| *<* 0.5) and moderate (0.5 *<* |*d*| *<* 0.8) effect sizes, respectively, according to Cohen’s d.

At the empirical level (Model-free Analysis), global brain network connectivity showed a general increasing trend from counting to early ACAM-J states, although these effects were modest and did not survive FDR correction (**Figure 3A**). Only the Executive Control and Default Mode networks exhibited uncorrected significant increases (Executive Control: t(15) = 2.32, p = 0.035; Default Mode: t(16) = 2.47, p = 0.025). In contrast, no significant differences were observed between early and late ACAM-J states, with only trend-level increases in Somato-Motor (t(18) = 2.02, p = 0.058) and Dorsal Attention networks (t(18) = 1.98, p = 0.063), indicating a relative stabilization of global brain network connectivity once the early phase is reached.

Consistent with this, whole-brain modeling revealed that the major shifts in dynamical regimes occur in the early ACAM-J (**Figure 3B**). When comparing counting to early ACAM-J states, the Default Mode and Visual networks showed moderate effect sizes (|d| > 0.5), while the Subcortical group and the Limbic network exhibited weaker effects (|d| > 0.2), indicating substantial movement toward the bifurcation point. In contrast, transitions from early to late ACAM-J were minimal, with only the Limbic network showing a weak effect size and all other networks remaining stable near the bifurcation point.

Together, these results suggest a progression in which large-scale reconfigurations are concentrated in the early ACAM-J states, followed by a relative stabilization within a near-critical regime in later states. This interpretation is supported by the full step-by-step analysis of transitions across ACAM-J states (**Figure S2**), which reveals that these early changes are not continuous but organized around discrete transition points corresponding to key meditative stages, including the initial engagement (J1), the peak of early ACAM-J (J4), and the transition into the late phase (J5), after which changes become comparatively smoother.

### Phenomenological changes and their relationship to DMN dynamics

To characterize the experiential profile across conditions, we analyzed phenomenological reports for stability of attention, width of attention, quality of ACAM-J, Jhana factor ratings, sights, sounds, physical sensations, and narrative thought stream (**Figure** 4). At the group level, the transition from the counting control condition into ACAM-J was associated with a marked reduction in narrative thought and a progressive broadening of attentional width, particularly from the earlier to later ACAM-J states. Physical sensations increased during the earlier ACAM-J states and decreased again in the later states, while reports of sights and sounds showed comparatively smaller changes. Stability of attention and the reported quality of ACAM-J remained consistently high across ACAM-J states relative to control. Jhana factor ratings were also high when the stage-specific factor was assessed, with the highest values observed for formlessness in the latest ACAM-J states (J7–J8).

**Figure 4:**
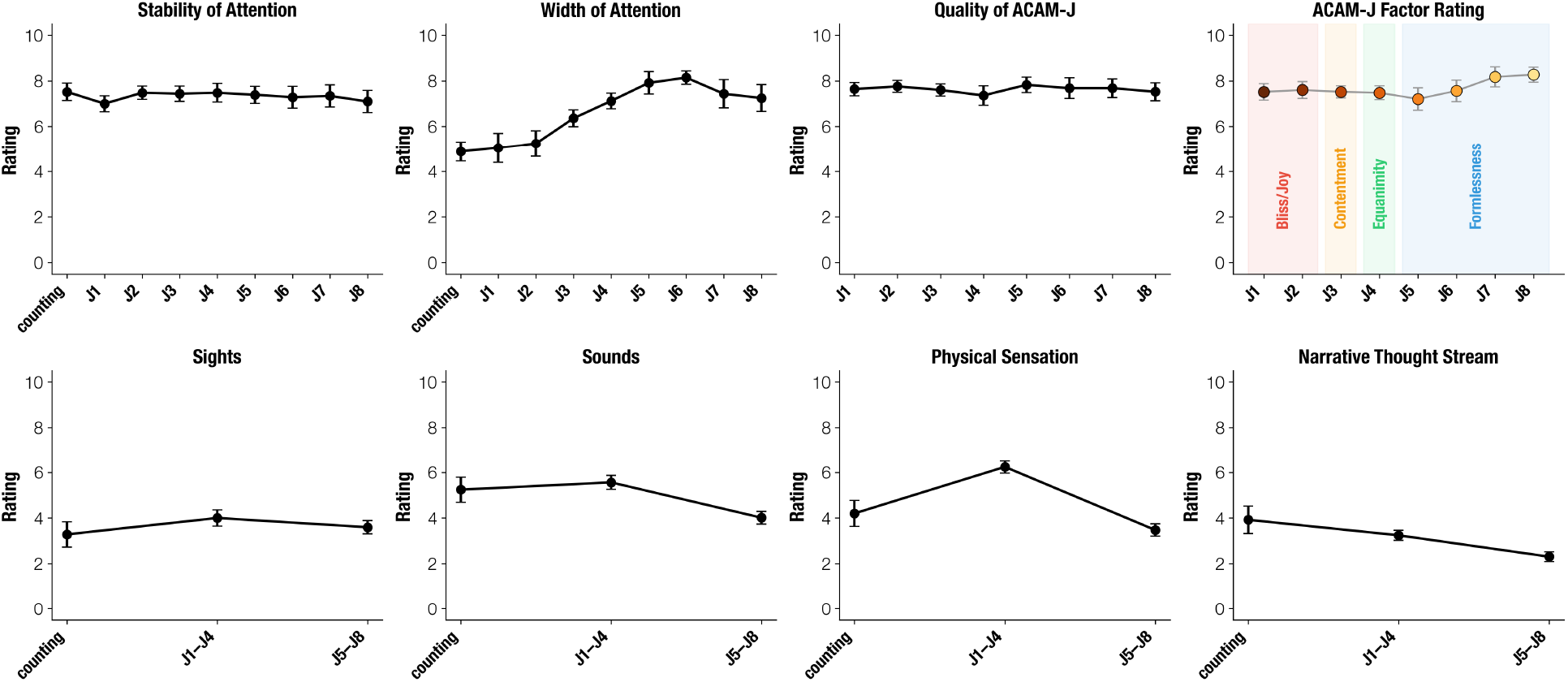
Phenomenological ratings across conditions. Group mean ratings (± SEM) for stability of attention, width of attention, overall ACAM-J quality, and stage-specific ACAM-J factor ratings are shown across the counting control and individual ACAM-J states (J1–J8). In addition, ratings for sights, sounds, physical sensation, and narrative thought stream are displayed for the counting condition and for the aggregate earlier (J1–J4) and later (J5–J8) ACAM-J states. All ratings were provided on a 0–10 scale, with higher values indicating greater intensity or clarity of the corresponding experiential dimension.

Given that the Default Mode Network exhibited the strongest dynamical shift in the model-based analysis, we next examined whether inter-individual variability in DMN distance to the bifurcation point tracked variability in phenomenological experience. Across subjects, the most robust associations emerged for narrative thought stream and width of attention (**Figure 5**).

**Figure 5:**
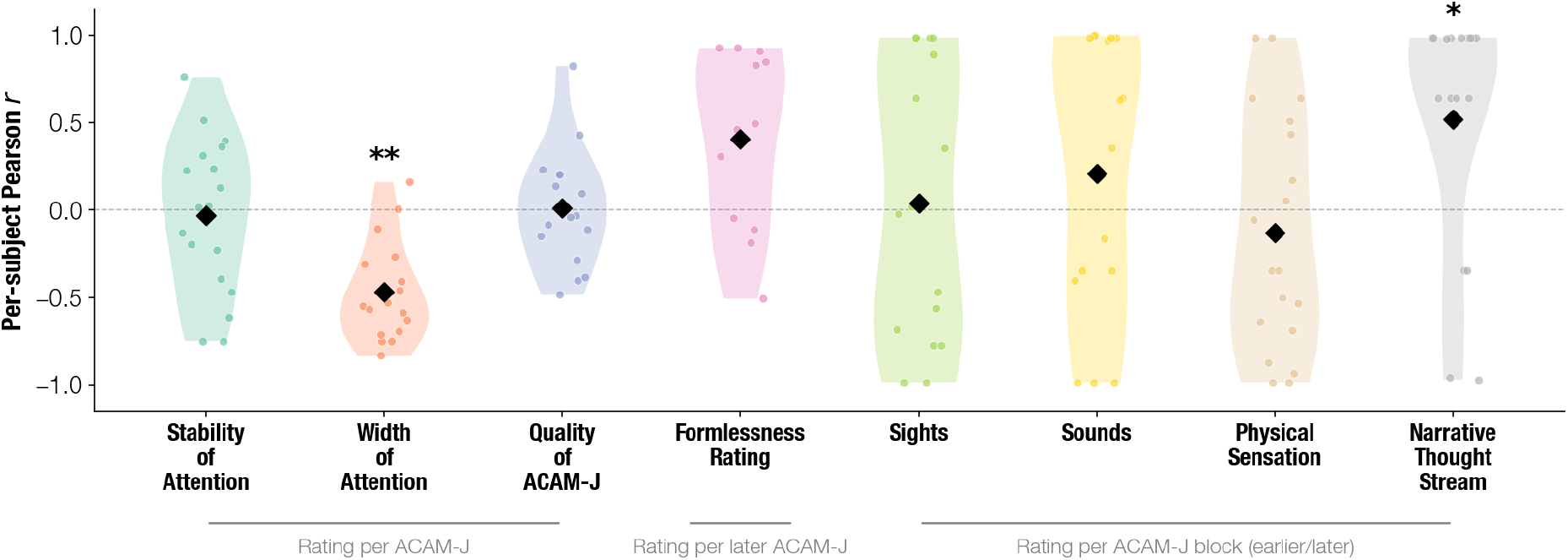
Subject-level associations between DMN dynamical regime and phenomenology. Pearson’s correlation coefficient (r) between DMN bifurcation parameter distance to bifurcation (|a|) and phenomenological ratings were computed within each participant across conditions. Violin plots display the distribution of subject-level correlation coefficients for each experiential dimension, with black diamonds indicating group means. Asterisks denote significant second-level effects after FDR correction across eight features (* *q* < 0.05; ** *q* < 0.01). Significant associations were observed for width of attention (*q* < 0.05) and narrative thought stream (*q* < 0.01).

Narrative thought showed a strong negative association with DMN distance to bifurcation (mean *r* = −0.47, FDR-corrected *q* < 0.01), indicating that conditions characterized by larger DMN distance values were associated with lower levels of reported narrative thought. This effect was highly consistent across individuals and survived both parametric and non-parametric second-level tests. Width of attention showed a positive association (mean *r* = +0.52, FDR-corrected *q* < 0.05), such that greater DMN distance to bifurcation point was associated with broader attentional scope. In contrast, stability of attention, quality of ACAM-J, Jhana factor ratings, sights, sounds and physical sensations showed no systematic relationship with DMN dynamics.

These findings were corroborated by a group-level analysis pooling all participant × condition observations. A linear mixed-effects model confirmed significant effects for width of attention (*β* = −134, FDR-corrected *q* < 0.001) and narrative thought stream (*β* = +80, *q* = 0.014), consistent with the subject-level results. In addition, formlessness ratings showed a significant positive association with DMN distance to bifurcation (*β* = +100, *q* = 0.014) that had been only borderline at the subject level (*q* = 0.050). The remaining features did not reach significance **(Figure S7)**. Together, these results link the model-derived dynamical reconfiguration of the DMN to specific dimensions of reported experience, most prominently the increase in width of attention and the reduction of narrative thought that characterizes the progression through ACAM-J.

## Discussion

Our central research goal was to identify mechanisms underlying advanced meditation and the construct of MPE, here studied in the context of ACAM-J. To address this, we first applied model-free analysis to describe the ACAM-J states. The empirical results showed a broad increase in large-scale functional integration during ACAM-J, particularly in the transition from counting to early ACAM-J states and in the deepest J7/J8 states used as empirical markers of MPE. Then using model-based analysis, we fitted whole-brain models to fMRI data across the eight ACAM-J states and examined dynamical regimes of resting-state networks, focusing in particular on the deepest ACAM-J states as empirical tests of MPE. Our analyses revealed that the gradient of RSN dynamical regimes observed in the counting control condition was markedly reduced in MPE, with most networks converging to near-critical dynamics. The DMN exhibited the largest shift, moving from a distant noise-driven regime during the control condition to near-critical dynamics through ACAM-J states. The trajectory across ACAM-J states was non-linear, with pronounced reconfigurations occurring at key meditative milestones, at J1, J4, and J5, followed by smoother changes during the later ACAM-J states. In what follows, we discuss the implications of these findings for the neuroscience of advanced meditation and MPE, related theoretical frameworks, and the science of consciousness. More generally, our results suggest that human flourishing may depend in part on the capacity of the brain to access flexible dynamical regimes without losing large-scale integration. Meditation may constitute one systematic method for modulating these dynamics endogenously

### RSN dynamics shift to criticality in ACAM-J

#### MPE as a near-critical working point

The healthy waking brain is thought to operate in an “optimal working point” near criticality, balancing sensitivity to perturbations with dynamical stability (Deco et al. 2012). In the context of whole-brain dynamical models, this notion of criticality refers to proximity to a dynamical instability (bifurcation), rather than to a thermodynamic second-order phase transition characterized by scale-free fluctuations and diverging correlation length. Empirical work in whole-brain modeling has demonstrated this by fitting a global bifurcation parameter (homogeneous across regions) to resting-state FC, showing it to lie near the bifurcation point (Deco et al. 2017). This is consistent with our genetic algorithm fitting results for the global bifurcation parameter in the counting condition, where a similar optimal working point is observed (**Figure S5**).

Crucially, however, we have seen that this global regime is sustained by a heterogeneous distribution of RSN and subcortical group dynamical regimes, with a few networks operating near the bifurcation point while most remain further away. Indeed, the whole-brain models in the counting condition showed an important gradation in the bifurcation parameter, with the Default Mode and the Visual network positioned at the extremes and furthest away from the bifurcation point (**Figure 3A**). In contrast, this observed gradient collapses for the later ACAM-J, which we use here as candidates for MPE. Networks that were previously far from the bifurcation point converged toward the near-critical regime (**Figure 3A**). Consistent with this dynamical convergence, the empirical analyses revealed a parallel increase in global brain network connectivity, with most RSNs and the subcortical group showing higher functional integration in MPE states compared to control. This suggests that during MPE, brain regions tend to operate close to criticality, with networks that were previously entrenched in their characteristic dynamics becoming more susceptible to the influence of the whole system (**Figure 6**). Interestingly, the global bifurcation parameter fitting results showed that the global working regime in MPE was also close to criticality (**Figure S5**). In this light, MPE appears to represent another optimal working point of the brain: one that is not sustained by heterogeneous dynamics across the functional gradient of RSNs and the subcortical group but by all RSNs and the subcortical group engaging in near-critical dynamics.

**Figure 6:**
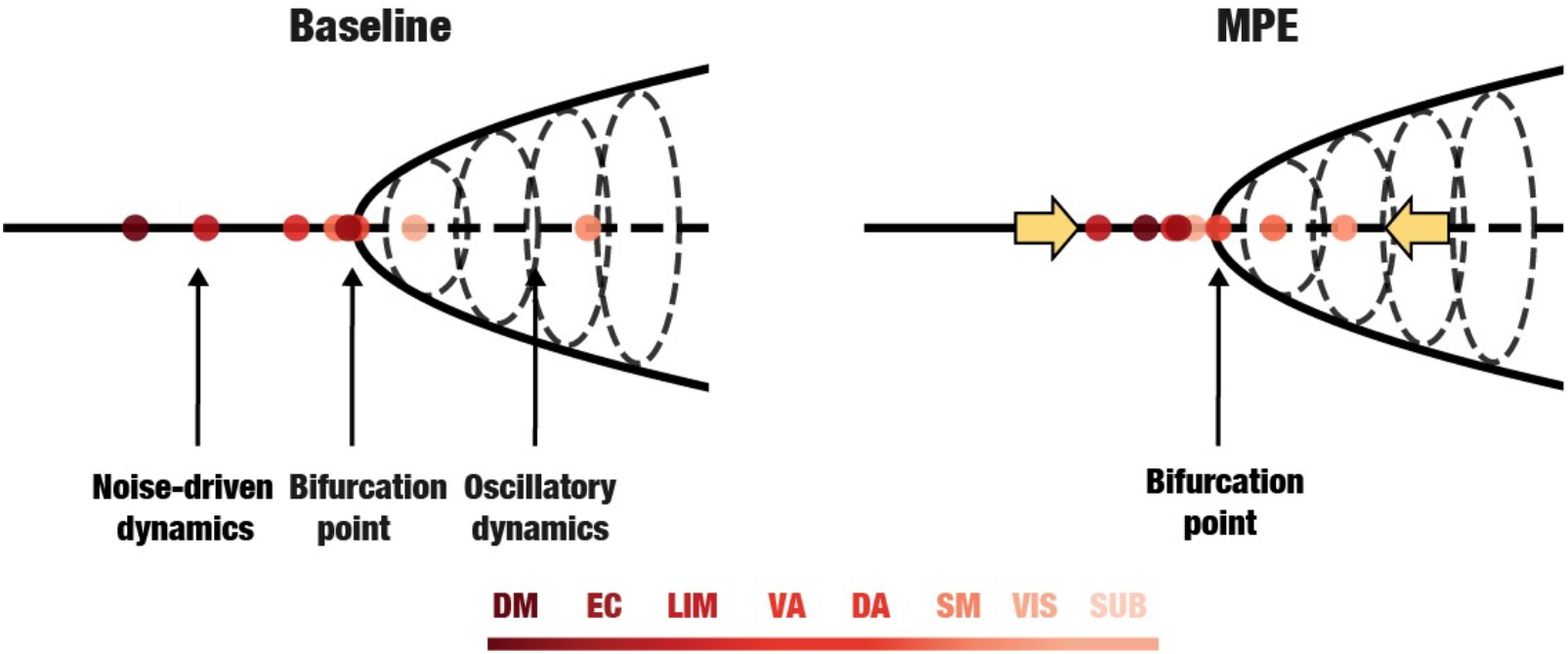
Brain dynamics shifts to criticality in MPE. The gradient from noise-driven to oscillatory dynamics observed in RSN bifurcation parameters during the counting condition (left) collapses in the later ACAM-J (right), with all networks shifting to critical dynamics. We hypothesize that, at baseline, the brain’s global regime is maintained through heterogeneity: different resting-state networks occupy distinct dynamical regimes, with the Default Mode and Visual network furthest from criticality. In contrast, during MPE, this heterogeneity collapses: all RSNs converge uniformly toward near-critical dynamics. This shift suggests that MPEs reflect an alternative optimal brain state, characterized not by differentiated dynamics across RSNs, but by a uniform engagement of all networks close to criticality.

The resulting global susceptibility of networks indicates a *latent* capacity to respond to perturbations. In dynamical terms, higher susceptibility implies that small inputs or fluctuations can more readily influence large-scale activity patterns, reflecting reduced constraint by stable attractors. In waking rest, RSNs are distributed across distinct dynamical regimes, supporting functional specialization, whereas in MPE they become uniformly susceptible, suggesting a shift to a more reconfigurable global regime. We interpret this increased reconfigurability as a form of higher or optimized, since information processing peaks under critical dynamics, flexibility, in which entrenched patterns of activity are temporarily relaxed. At the experiential level, such a regime may be described as one of maximal “openness”, in which habitual modes of perception and self-related processing are softened. This dynamical flexibility may allow the brain to reset or reorganize its dynamical repertoire, including deeply rooted attractors, helping to explain why emerging from these states is often associated with marked shifts in perception, or *insights*, which are considered in contemplative traditions as central to the transformative potential of jhāna practice (Burbea 2014; Tulver et al. 2023).

#### Increased DMN susceptibility in MPE

Beyond the global convergence to criticality, our results revealed network-specific shifts in dynamical susceptibility. Networks closer to the bifurcation point are more sensitive to perturbations and therefore more strongly shaped by, and responsive to, large-scale system dynamics. The DMN showed the largest change in the bifurcation parameter. In the control condition, the DMN was the furthest from the bifurcation point, and in MPE it became markedly closer to it, becoming more susceptible to perturbation. Importantly, this increased susceptibility reflects reduced local damping within the fitted dynamical model, rather than intrinsic scale-free activity of the DMN. This indicates that its dynamics are no longer dominated by intrinsic activity but instead are more responsive to the system as a whole. Such a shift suggests an important role of DMN in the large-scale reorganization of brain activity during later ACAM-J states, and may reflect a transition from within-to between-network integration (Treves et al. 2025). This interpretation is consistent with recent proposals that position the DMN as a central hub mediating the convergence and divergence of large-scale brain dynamics (Luppi et al. 2025), as well as empirical reports of altered DMN activity in meditation (Garrison et al. 2015), and more specifically in ACAM-J (Ganesan et al. 2024).

More broadly, the DMN has been consistently implicated in self-referential cognition, autobiographical memory, and narrative forms of selfhood (Buckner et al. 2008; Qin et al. 2011). From this perspective, the increased susceptibility of the DMN during MPE suggests a qualitative shift in self-processing. By moving closer to the critical point, DMN dynamics may become less rigid and more adaptable, allowing self-related processes to be more responsive to ongoing global brain dynamics rather than being dominated by internally driven activity. Such increased flexibility is consistent with phenomenological reports of attenuated identification and reduced fixation on self-related content in minimal phenomenal experience (Metzinger 2024), and with contemplative accounts that emphasize the cultivation of a more Eluid and less entrenched mode of self-experience (Dahl et al. 2015).

Importantly, our subject-level analyses provide empirical support for this interpretation. Across individuals, decreased DMN distance to bifurcation point was robustly associated with reduced narrative thought and increased width of attention (Figure 5). These effects were highly consistent across participants and were not observed for stability of attention, formlessness factor ratings, or other type of experiential content. This specificity suggests that DMN reconfiguration is selectively linked to the attenuation of narrative self-processing and the expansion of attentional scope, rather than reflecting a global intensification of meditative experience. This is consistent with clinical findings linking DMN hyperactivity and reduced integration to psychopathologies associated with rumination and self-referential thought, such as depression or anxiety (Hamilton et al. 2015; Sylvester et al. 2012).

### ACAM-J criticality and related theoretical accounts

These findings resonate with recent computational work within the free energy principle framework, which models MPE in terms of variational free energy minimization and likewise emphasizes its relation to critical dynamics (Sandved-Smith 2024). In addition, recent mechanistic accounts frame ACAM-J as a process of flattening hierarchical attractors, which manifests phenomenologically as a progressive *defabrication* of experience (Lopez-Sola et al. 2025). Flattening attractors in this way implies a dynamical regime where existing patterns of activity lose stability, and the system as a whole becomes more susceptible to perturbation (Laukkonen & Slagter, 2021). This description aligns closely with our finding that RSN dynamics converge toward near-critical dynamics and higher susceptibility in the later ACAM-J.

We also described this process as a sequential disruption of the hierarchical models, which is phenomenologically described as a progressive “letting go” of increasingly ‘basic’ experiences (space, consciousness, nothingness, …) (Burbea 2014; Prest et al. 2024; Laukkonen & Slagter, 2021). Increasingly, each state involves the release of rigid priors, yielding greater sensitivity and reduced constraint, consistent with the approximation to criticality that we observe here. MPE may represent a theoretical limit of this spectrum of letting go, and our results characterize a series of empirically accessible meditative states whose dynamics progressively approach this regime as brain dynamics come closest to the critical point.

Relatedly, recent studies using the active inference framework have proposed that the progression through the ACAM-J states and towards MPE may be driven by a systematic reduction of prior precision across the hierarchy (Laukkonen et al. 2021; 2025; Mago et al. 2024; Prest et al. 2024; Sandved-Smith et al. 2024; Tal et al. 2025), also aligning with the description of a progressive flattening of hierarchical attractors. Moreover, lower precision would make the generative model more malleable, enabling the reconstruction of priors in new ways, which resonates with the transformational aspect mentioned above.

Converging evidence for the prospect of MPE as near-critical states comes from recent studies on meditative practices and psychedelics using measures that reflect various signatures of criticality. A recent study found that ACAM-J alters functional connectivity gradients, collapsing the usual hierarchical separation between sensory and higher-order regions and promoting a more globally integrated mode (Demir et al. 2025). Such a collapse of hierarchical differentiation is consistent with our finding that MPE involves a convergence of RSN dynamics toward criticality. Additional support comes from EEG criticality studies of meditation-induced cessation events (van Lutterveld et al. 2025). These cessation events often mark the fruition of insight practice and involve profound transformations in awareness, and the fact that they also exhibit signatures of criticality strengthens the view that such dynamics may be a common marker of MPE.

A further line of evidence relates to entropy and complexity. Dynamical critical regimes are often characterized by increased complexity (Chialvo 2010, Ruffini 2023, 2024, 2025), suggesting that entropy and complexity should increase progressively across the ACAM-J states and peak during MPE. Such a hypothesis was explicitly advanced in recent computational work (Mago et al. 2024), and although higher complexity and entropy have been reported in meditative states more broadly (Atad et al. 2025), specific evidence for ACAM-J is still lacking.

In addition to the canonical resting-state networks, our results indicate that the subcortical system also moved toward a near-critical regime. This suggests that deeper structures, including those involved in arousal regulation, salience gating, and homeostatic control, may participate in the same global shift toward increased susceptibility. These structures are known to modulate cortical gain and influence the overall stability of hierarchical models. Their approach toward the critical regime may therefore reflect a coordinated loosening of global constraints that supports the phenomenological transition toward meditative absorption states. This complements the attractor flattening account, since subcortical drive and gain modulation likely play an important role in enabling cortical models to lose stability in a controlled way (Friston 2023).

Finally, the mechanistic account of MPE we have discussed resonates with Metzinger’s interpretation of MPE as the experience of an *epistemic space* (Metzinger 2024), an “inner space holding a very large number of possibilities for knowing the world and ourselves”. It may also be compatible with Metzinger’s proposal that MPE corresponds to a representation of *tonic alertness* (Metzinger 2020). In our view, the collapse of heterogeneous RSN dynamics into a uniformly near-critical regime reflects a stripping away of hierarchical models, and what remains could be this minimal, non-egoic alertness. Accordingly, when object perception, self-representation, and executive control are released, the brain may settle into a globally susceptible dynamics whose neural signal corresponds to this “alertness Eield”. This interpretation is further supported by the fact that subcortical systems also shift toward the near-critical regime, which suggests that they may help maintain this minimal background of wakeful presence.

#### Relationship to meditative absorption and non-dual awareness

It is important to note that our proposal of maximal susceptibility in MPE is compatible with deep meditative absorption states (Sparby et al. 2024), in which external objects are not actively represented. We hypothesize that in these states the absence of active tracking of sensory objects frees the system from the usual exogenous constraints while leaving the dynamics open. The system remains highly critical in the sense of being fully available for reconfiguration, which is precisely what is often observed upon emerging from these states.

But such absorption states are clearly not compatible with everyday functioning, given the lack of representational content. A natural question, then, is whether there are modes of experience that are maximally susceptible yet compatible with ordinary activity. A promising candidate is *nondual awareness*, which can co-occur with phenomenal content, as described in many of Metzinger’s reports (Metzinger 2024). In our recent work, we suggested that nondual awareness is not a *minimal* state, but is related to some MPEs in that awareness itself comes to the foreground of experience (Lopez-Sola et al. 2025). We proposed that this may involve an *opacification* of the modeling hierarchy, creating a distinct form of global flexibility. A provocative hypothesis is that nondual awareness also reflects a mode of near-critical brain dynamics and maximal openness, but one that remains compatible with representing other forms of phenomenal content (as opposed to deep absorption states). Testing this possibility would require applying the present modeling framework to empirical data on nondual states.

### The progression through ACAM-J

Results of the current study suggest that the collapse of RSN dynamics toward criticality is non-linear. Most networks approached near-critical dynamics within the early ACAM-J states and remained close to criticality during the transition through the later ACAM-J states. At a finer scale, however, each RSN followed a distinct trajectory (Figure S2).

Notably, the most dramatic changes coincided with phenomenological milestones of ACAM-J (Brasington 2015; Yang et al. 2024). The first was the entry into J1, which is described as a deep absorption state characterized by intense bliss (Sparby et al. 2024). Given that this transition involves moving from a control task (counting) into meditation, a major reconfiguration in brain dynamics is to be expected, and indeed we observed such a shift. The second occurred at J4, the peak of the *early* ACAM-J, dominated by equanimity (Sparby et al. 2024). The third was the passage from *early* to *late* ACAM-J (J4–J5), where the perception of form (sights, sounds, body sensations) is thought to dissolve, leaving only “boundless space” (Brasington 2015; Burbea 2014).

At each of these milestones, some of the largest changes were observed in the Default Mode and Visual networks. This pattern is consistent with their position in the control condition (furthest from the bifurcation), and with our finding that the DMN undergoes the strongest overall reconfiguration. The pronounced involvement of the Visual network in the transition to the later ACAM-J may relate to the reported loss of object perception, although this interpretation should be treated cautiously given that visual network dynamics also differ between control conditions.

By contrast, the later ACAM-J were not marked by large, discrete changes, and most networks remained stably near the bifurcation (see Supplementary Figure S2). The absence of large reconfigurations in the later ACAM-J may reflect their phenomenological character as more pacified states, in which transitions are experienced as slower and more gradual (Brasington 2015). This pattern is also consistent with recent theoretical accounts of ACAM-J and MPE (Lopez-Sola et al. 2025), in which the progression through the later ACAM-J corresponds to the flattening of increasingly deep and subtle attractors. In this view, the later ACAM-J consolidate the system into a broadly flattened dynamical landscape, with residual network heterogeneity, rather than driving further large-scale reconfigurations.

This nonlinear collapse of network dynamics toward near-critical regimes may reflect how the brain endogenously accesses MPE through meditation. The transition toward a globally susceptible state may be constrained by biological dynamics, such that networks cannot shift linearly toward criticality together but must do so in stages. Whether other pathways to MPE, such as psychedelic states, follow a similar stepwise progression remains an open question.

### Implications for consciousness research

The interest for MPE was originally motivated by the search for a minimal model of consciousness, providing a framework to study what the most basic form of conscious awareness might look like (Metzinger 2020). Recent proposals argue that advanced meditation offers a promising experimental strategy toward such a minimal model, by systematically reducing experiential content while maintaining abstract forms of awareness (Lieberman et al. 2025). In this context, deep ACAM-J states constitute particularly strong candidates for MPE. Our findings suggest that these states may correspond to a global collapse of large-scale differentiation into a near-critical regime marked by maximal susceptibility and extended spatio-temporal correlations. This carries several implications for theories of consciousness.

First, the convergence of RSN dynamics toward criticality in MPE is consistent with theories that emphasize global accessibility and integration as core features of conscious states. For instance, the Global Workspace Theory proposes that consciousness arises from widespread ignition or broadcast across the brain (Baars 2005). A uniformly near-critical regime would naturally support such ignition, since the entire system is highly susceptible to perturbations. In this light, MPE could be understood as a configuration with maximal *capacity* for broadcast, even if little or nothing is actually broadcast. Similarly, Integrated Information Theory highlights integration as a hallmark of consciousness (Tononi 2008); our results suggest that MPE may represent a configuration in which integration is achieved not through large-scale specialization of RSNs but through increased spatio-temporal correlation across networks.

Second, the global collapse we observed during MPE offers an interesting perspective on its status as a minimal model. If consciousness in its most stripped-down form is associated with undifferentiated, maximally near-critical dynamics, this may point toward a primordial mode of conscious functioning. From a developmental and evolutionary perspective, early forms of consciousness in fetuses, infants, or non-human animals might resemble such globally flexible, weakly differentiated states, before rigid hierarchical control is established. Evidence for this comes from developmental neuroimaging, which shows that functional connectivity in fetuses and newborns is dominated by global, low-dimensional patterns, with higher-order network differentiation only emerging later in infancy and childhood (Gao et al. 2017). Comparative research also suggests that simpler organisms and non-human animals often display globally integrated architectures, lacking the strong hierarchical segregation characteristic of adult human brains (Van den Heuvel et al. 2016).

This perspective highlights the transformative potential of MPE. In ordinary functioning, hierarchies and rigid dynamics constrain the system, ensuring survival and goal-directed behavior but limiting flexibility. Through practices such as ACAM-J, these constraints can be temporarily released, allowing the system to return to a globally open regime. Such reversals may explain why some minimal experiences, such as what has been called “pure awareness,” are so often described as transformative. This framework thus provides a robust foundation for understanding meditative endpoints more broadly (Sparby and Sacchet, 2025).

### Limitations and future research

We chose to parametrize the whole-brain models in terms of resting-state networks. This was a natural first step, given their robustness and frequent use in both empirical and modeling studies. However, alternative priors could be equally or more informative. For example, functional gradients might capture large-scale cortical organization, and neurotransmitter receptor maps could link model parameters more directly to neuromodulatory dynamics. Additionally, in our study subcortical regions were modeled as a single group, despite their functional heterogeneity, so the observed shift toward near-critical dynamics may be driven by specific nuclei, such as the thalamus or caudate, rather than reflecting a uniform subcortical effect, though the supplementary network analysis indicated the subcortical functional shifts to be more of a global effect (Figure S8). Future work should explore these alternative parametrizations, as they may reveal additional insights into the mechanisms of advanced meditation practices such as ACAM-J. Lastly, the divergence between control conditions highlighted that not all baseline tasks provide equivalent reference points. Differences observed between ACAM-J and control conditions should therefore be interpreted in relation to the specific baseline employed, rather than as absolute markers of meditative states.

## Conclusions

In conclusion, our study provides unique insights into the neuroscience of advanced meditation, ACAM-J, MPE, and ‘pure awareness’. By drawing on advanced meditation we have established modeling and analyses of functional data to understand underlying neural and psychological mechanisms and dynamical regimes. In doing so, this study provides the type of formal modeling that is needed to powerfully bridge first-person phenomenology with third-person neural data, relating formal mechanisms to testable neurobiological hypotheses, while avoiding pitfalls of phenomenologically-inspired “mathematical storytelling”. As the first whole-brain modeling study of advanced meditation, ACAM-J, and MPE, this study contributes to a foundation for a neuroscientific mechanistic understanding of advanced meditation, and to inspiring further integration of meditative phenomenology with computational neuroscience.

## Contributions

J.V. and E.L-S. conceived the modeling approach and study reported here. M.D.S. conceived of and funded the broader study. J.V. and E.L-S. designed and performed the analyses, with methodological input from Y.S-P., M.L.K., G.R. and G.D.. W.F.Z.Y, T.S., R.P. and M.D.S. collected the meditation dataset. W.F.Z.Y. handled and preprocessed the fMRI data. J.V., Y.S-P, M.L.K and G.D. developed and implemented the whole-brain modeling framework, including optimization procedures. M.D.S. and G.D. provided conceptual guidance and supervised the work. R.E.L assisted with conceptual guidance, interpretation of results, narrative framing, and figures. J.V. and E.L-S. wrote the first draft of the manuscript. All authors contributed to the interpretation of results, manuscript revisions, and approved the final version.

## Supplementary Information

### Supplementary Methods

#### Neuroimaging Acquisition and Preprocessing

MRI data acquisition parameters followed the original study protocol (Yang et al. 2025b): high-resolution 7T fMRI (SIEMENS MAGNETOM Terra, 32-channel head coil) with whole-brain coverage, along with structural T1-weighted scans. Physiological signals (heart rate, respiration) were recorded concurrently to allow correction for physiological noise.

Data preprocessing steps replicated those from the single-case design, including de-spiking and physiological noise regression (RETROICOR), slice-timing and distortion correction, motion correction and scrubbing, normalization to standard space, band-pass filtering (0.01–0.1 Hz), spatial smoothing (2mm FWHM), and z-scoring of time series. Each run was segmented into ACAM-J states (J1–J8) or control-task periods, yielding condition-specific time series for each subject.

#### Structural Connectivity

The *N*×*N* matrices of structural connectivity (*C*) used for the network model were derived from a probabilistic tractography-based normative connectome provided as part of the leadDBS toolbox (Horn et al. 2015). This normative connectome was generated from diffusion-weighted and T2-weighted Magnetic Resonance Imaging (MRI) from 32 healthy participants (mean age 31.5 years, 14 females) from the Human Connectome Project (HCP). The diffusion-weighted MRI data were recorded for 89 minutes on a specially designed MRI scanner with more powerful gradients than conventional scanners. The dataset and acquisition protocol details are available in the Image & Data Archive under the HCP project (https://ida.loni.usc.edu/). DSI Studio (Yeh 2025) was used to implement a generalized q-sampling imaging algorithm on the diffusion data. A white matter mask, derived from the segmentation of the T2-weighted anatomical images, was used to co-register the images to the b0 image of the diffusion data using the SPM12 toolbox (Ashburner et al. 2014). Within the white-matter mask, 200,000 most probable fibres were sampled for each participant. The fibres were then transformed to the standard Montreal Neurological Institute (MNI) space using a nonlinear deformation field derived from the T2-weighted images via a diffeomorphic registration algorithm. The individual tractograms were aggregated into a joint dataset in MNI standard space, resulting in a normative tractogram representative of a healthy young adult population and made available in the leadDBS toolbox. The *N* ×*N* matrices were computed from the normative tractogram using the Schaefer-200 parcellation scheme (Schaefer et al. 2018) with *N* = 200 cortical parcels and Tian Scale 2 parcellation scheme with *N* = 34 subcortical parcels (Tian et al. 2020). Structural connectivity values *C*(*n,p*) were calculated as the number of tracts between each pair of brain areas *n* and *p*.

#### Genetic algorithm parameters

Each optimization run began with a population of 10 individuals with random bifurcation parameter values close to zero (centered around *a* = −0.01) and progressed through cycles of elite retention, crossover, and mutation until convergence criteria were met (maximum 20 generations or stabilization of the best solution). In each generation, 20% of individuals were preserved through elite selection, 60% were produced by crossover of parent solutions, and 20% by mutation. Forty independent runs were performed for each ACAM-J state, and the resulting distributions of parameters were used to assess stability.

## Supplementary results

### Analyzing the later ACAM-J states as candidate MPE states

**Figure S1:**
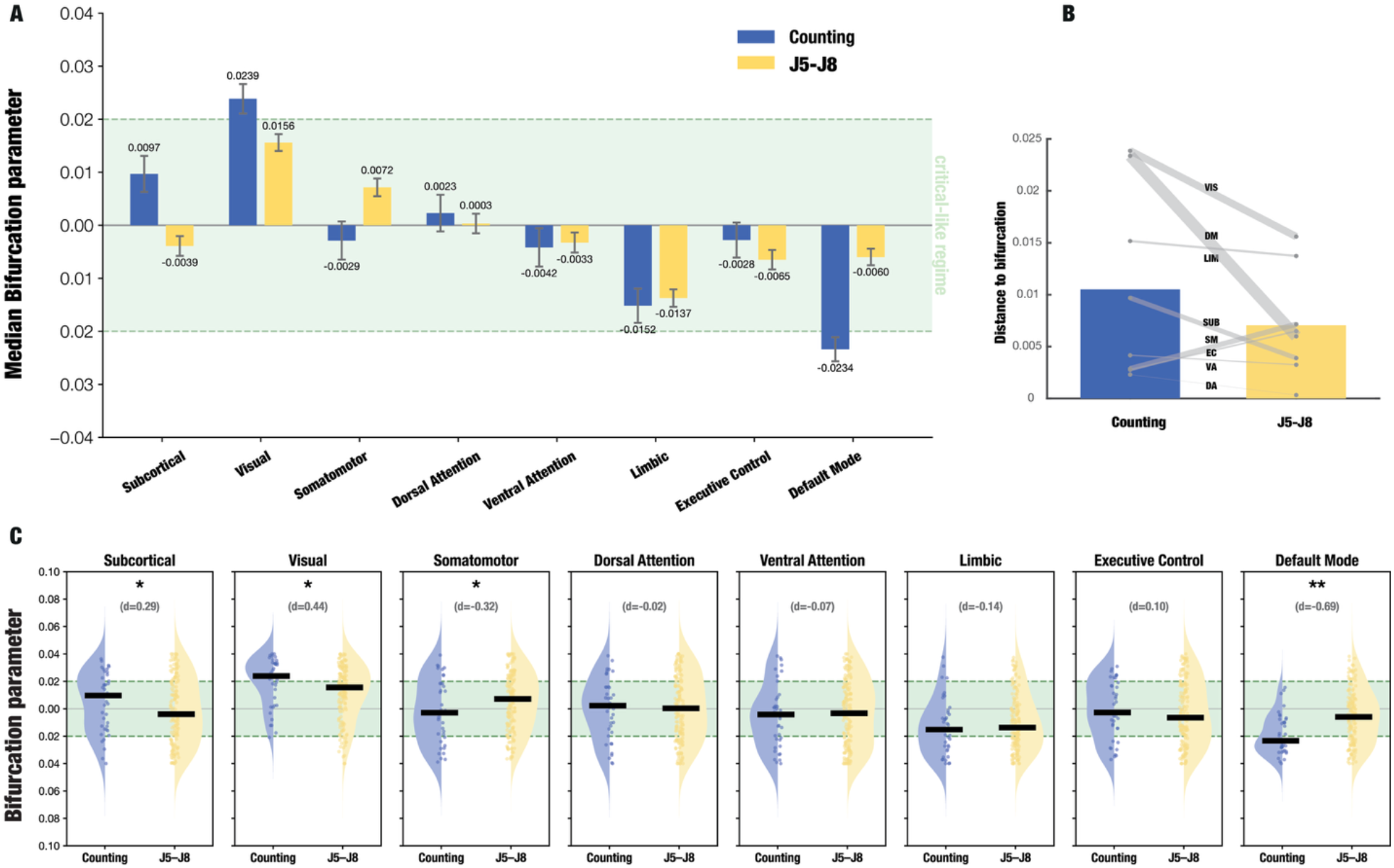
ACAM-J shifts network dynamical regimes to near-critical dynamics. Bifurcation parameter values per RSN in the counting condition and the later ACAM-J states (J5-J8, the *formless* states). **A**: Median bifurcation parameter values per RSN in the counting and *formless* conditions. The green shaded area marks the near-critical region (|*a*| *<* 0.02). **B**: Median distance of RSNs to the bifurcation point (*a* = 0). Dots indicate median distances per RSN, and line width represents the effect size according to Cohen’s d. **C**: Distribution of bifurcation parameters per RSN for forty runs of the genetic algorithm in each condition. Solid lines indicate median values, * and ** indicate weak (0.2 *<* |*d*| *<* 0.5) and moderate (0.5 *<* |*d*| *<* 0.8) effect sizes according to Cohen’s d.

### Mechanistic progression through ACAM-J states

**Figure S2:**
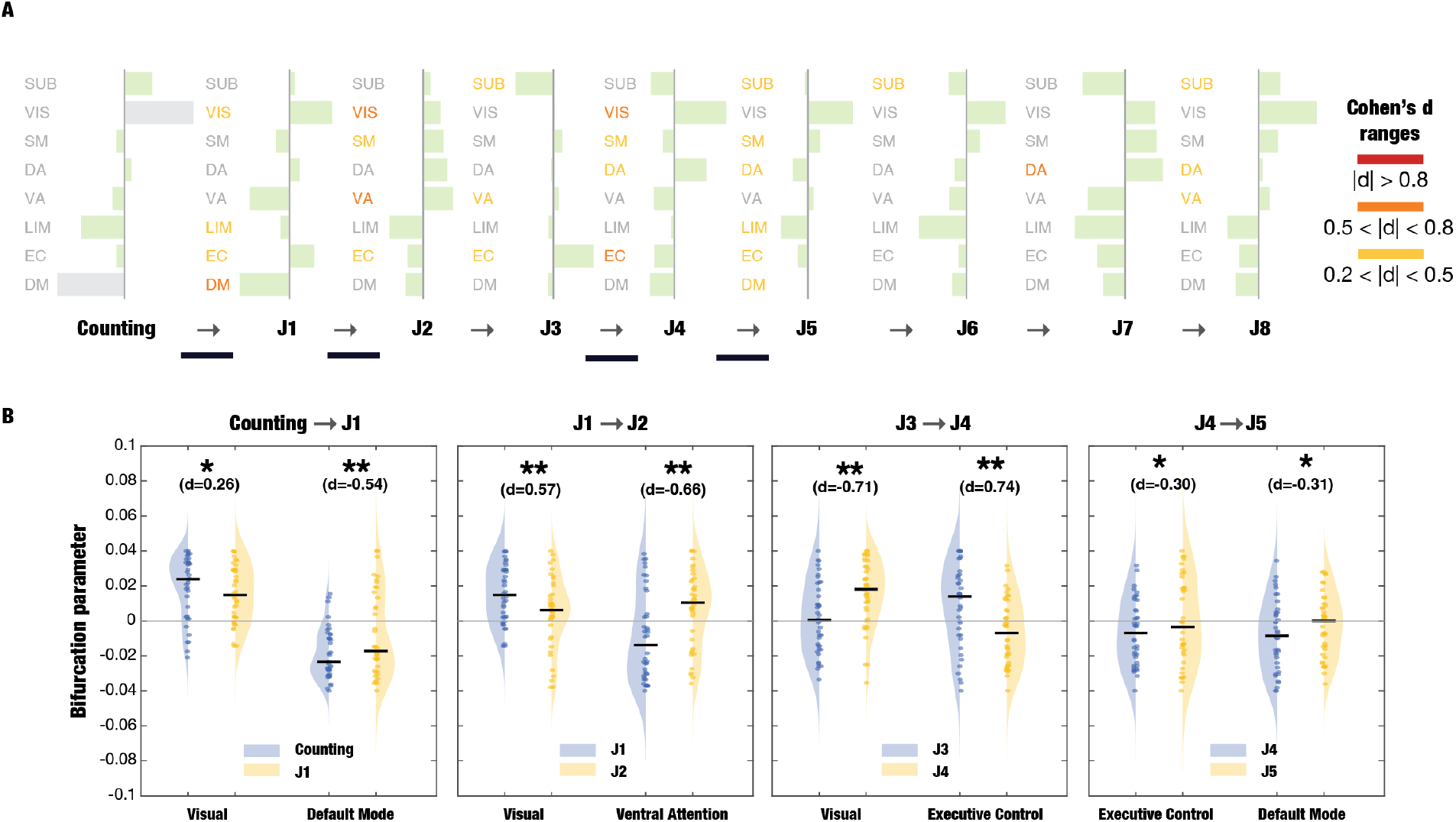
Mechanistic progression through ACAM-J. **A:** Progression of the bifurcation parameter in each RSN across the different ACAM-J states. The trajectory through ACAM-J is marked by key transition points (solid black line) in the transitions to J1, J4, and J5, after which the changes in dynamics are smoother. Bars indicate median values, green bars indicate proximity to bifurcation point (|*a*| *<*0.02). Colors indicate the Cohen’s d effect size in the change from the previous condition. **B:** Distribution of bifurcation parameters for the Visual and Default Mode networks in the three key transitions. Solid lines indicate median values, * and ** indicate small (|*d*| *>* 0.2) and moderate (0.5 *<* |*d*| *<* 0.8) effect sizes according to Cohen’s d.

### MPE state (J7-8) comparison to ACAM-J1 and ACAM-J4

**Figure S3:**
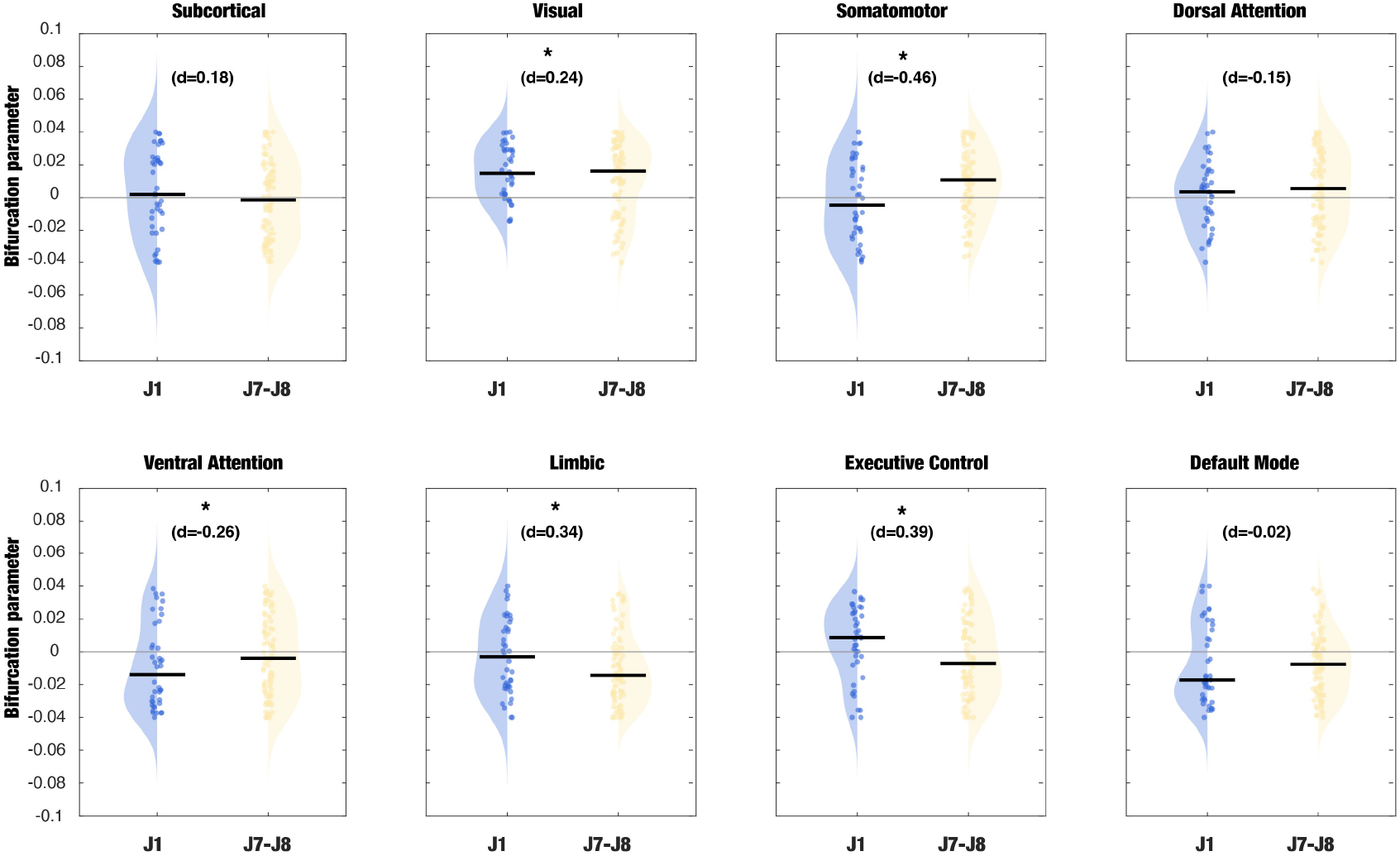
Comparison of the bifurcation parameter between early absorption (ACAM-J1) and late absorption (ACAM-J7/J8) across networks (see below).

**Figure S4:**
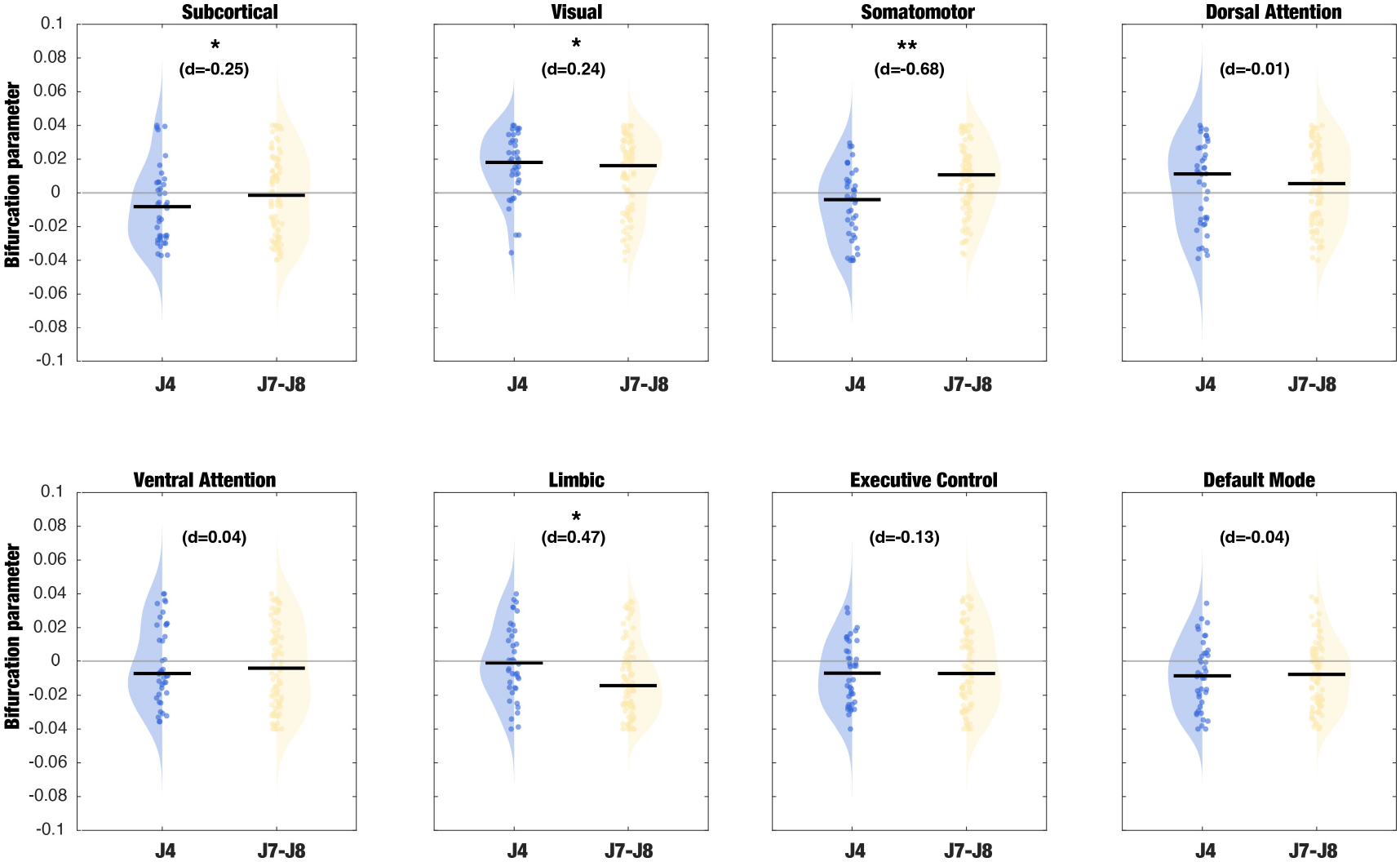
Comparison of the bifurcation parameter between the peak of the early ACAM-J (J4) and the last ACAM-J states (J7/J8) across networks. Violin plots display subject-level distributions, with dots representing individual participants. Solid lines indicate median values, * and ** indicate weak (0.2 *<* |*d*| *<* 0.5) and moderate (0.5 *<* |*d*| *<* 0.8) effect sizes according to Cohen’s d.

### Global parameter whole-brain fitting

**Figure S5:**
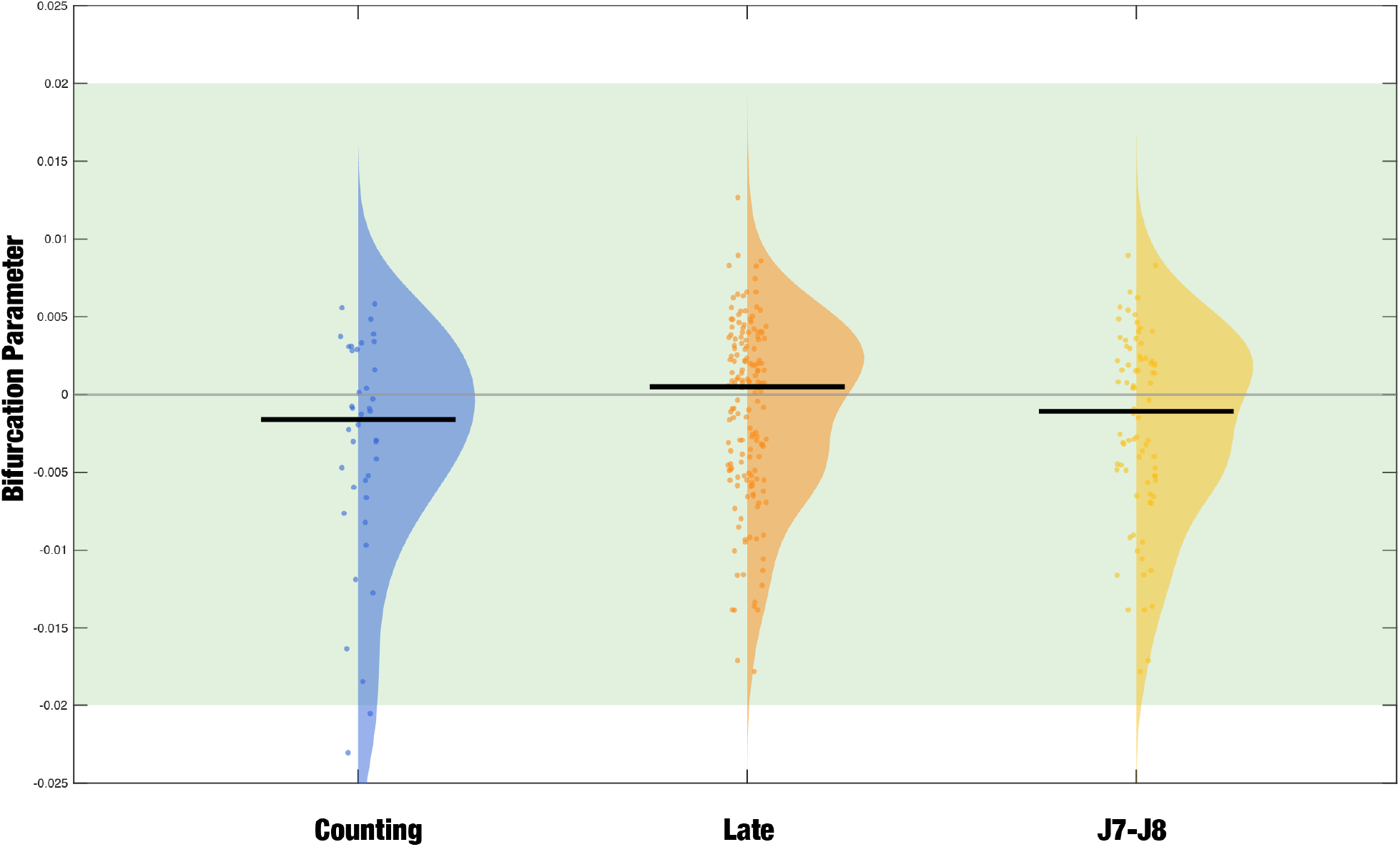
Results of the global parameter whole-brain model fitting. We optimized only one bifurcation parameter to fit the models, instead of parametrizing the bifurcation parameter of each RSN. We show the distribution of bifurcation parameter values for the counting, late ACAM-J (J5-J8) and J7-J8 conditions in forty iterations of the genetic algorithm. The bifurcation parameter remained close to the bifurcation (near-critical regime) in all the control conditions and ACAM-J states.

### Comparing the counting and memory control condition

**Figure S6:**
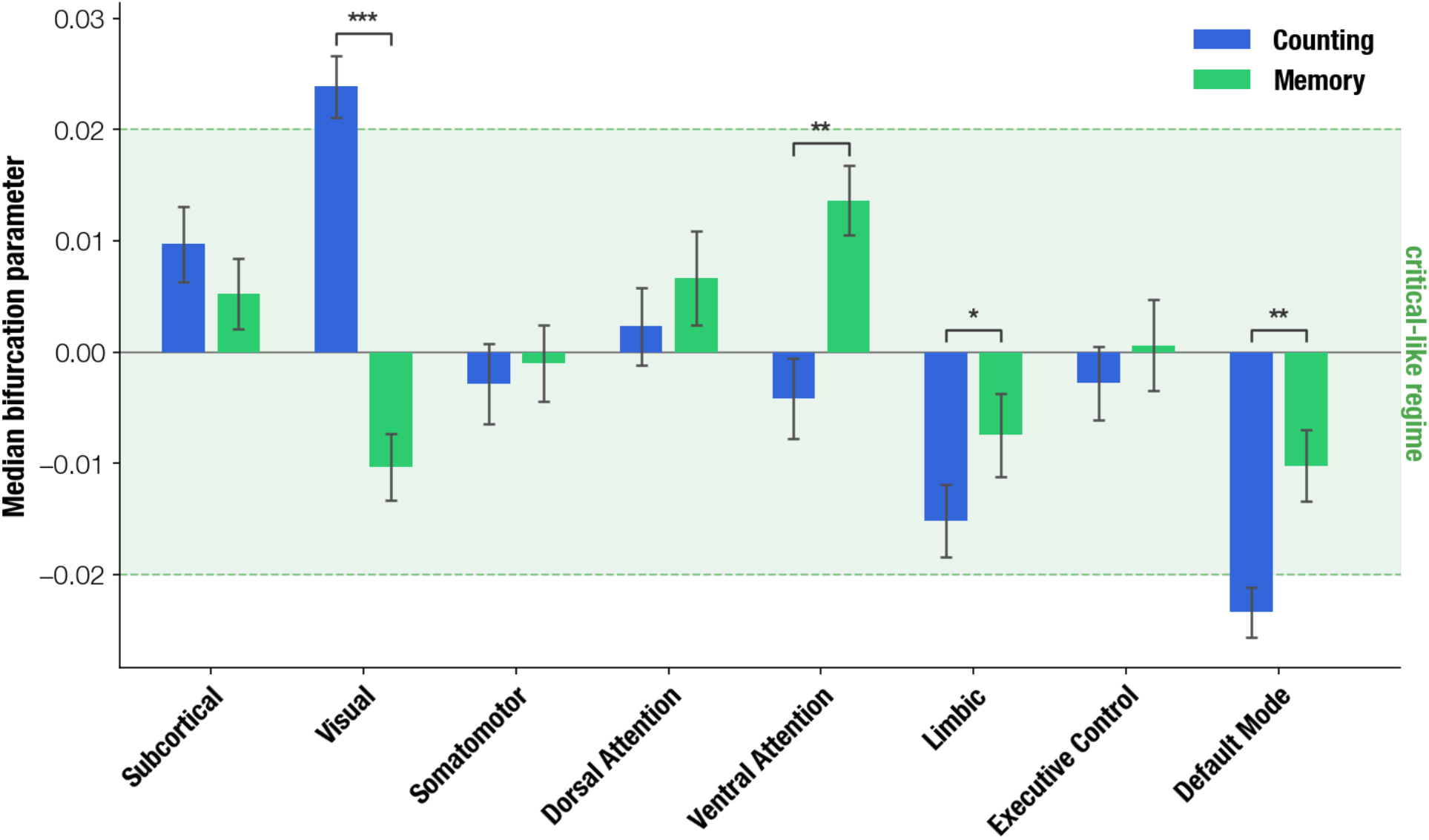
Dynamical regimes in counting and memory control conditions. Bifurcation parameter values per resting-state network (RSN) for the counting and memory control conditions. The green shaded area indicates the near-critical regime (|a| < 0.02). Bars represent median values across optimization runs, with error bars indicating variability across runs. Asterisks denote effect sizes based on Cohen’s d (* weak, ** moderate). The two control conditions exhibit systematic differences across RSNs. The counting condition reproduces the canonical resting-state pattern reported in prior whole-brain modelling studies, with higher-order networks in subcritical regimes and sensory networks in supercritical regimes. In contrast, the memory condition shows shifts consistent with increased internally generated processing, particularly in visual and attention networks. These results indicate that the two control conditions instantiate distinct dynamical baselines rather than interchangeable references.

### Correlations between phenomenology and DMN distance to bifurcation

**Figure S7:**
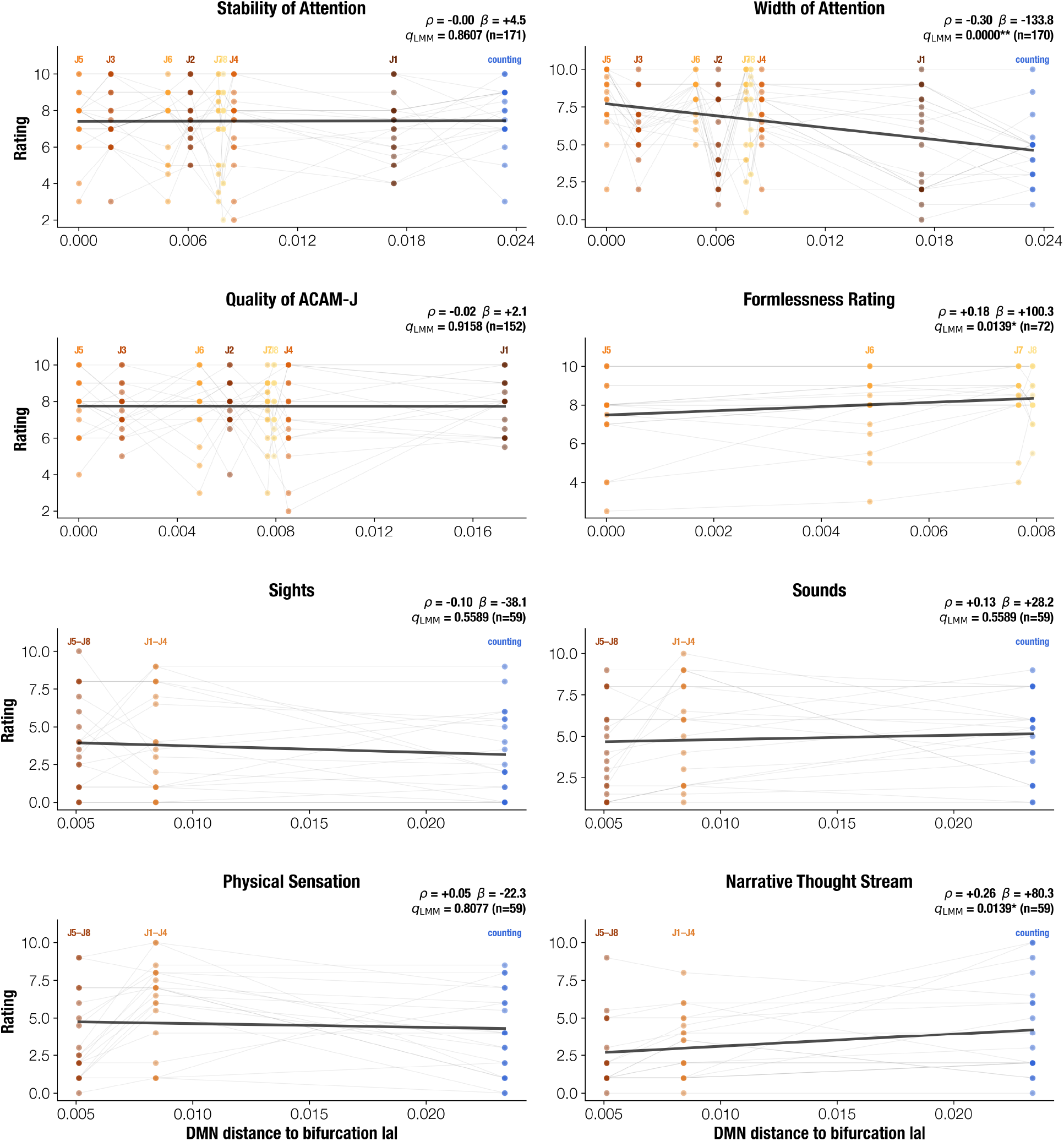
Correlations between phenomenology and DMN distance to bifurcation. Each panel shows one phenomenological feature, with DMN distance to bifurcation (x-axis) plotted against phenomenological ratings (y-axis). Each dot represents one participant in one condition, colored by condition. Thin grey lines trace individual participant trajectories across conditions. The black line shows the linear fit across all observations. For content-related dimensions (Sights, Sounds, Physical Sensation, Narrative Thought Stream), earlier (J1–J4) and later (J5–J8) ACAM-J states are shown as aggregate levels. For each panel, *ρ* is the Spearman rank correlation, *β* is the fixed-effect slope from a linear mixed-effects model *(rating ∼ bifurcation + (1* | *participant)*), and *q*_LMM_ is the corresponding FDR-corrected p-value. **q < 0.01, *q < 0.05.

**Figure S8:**
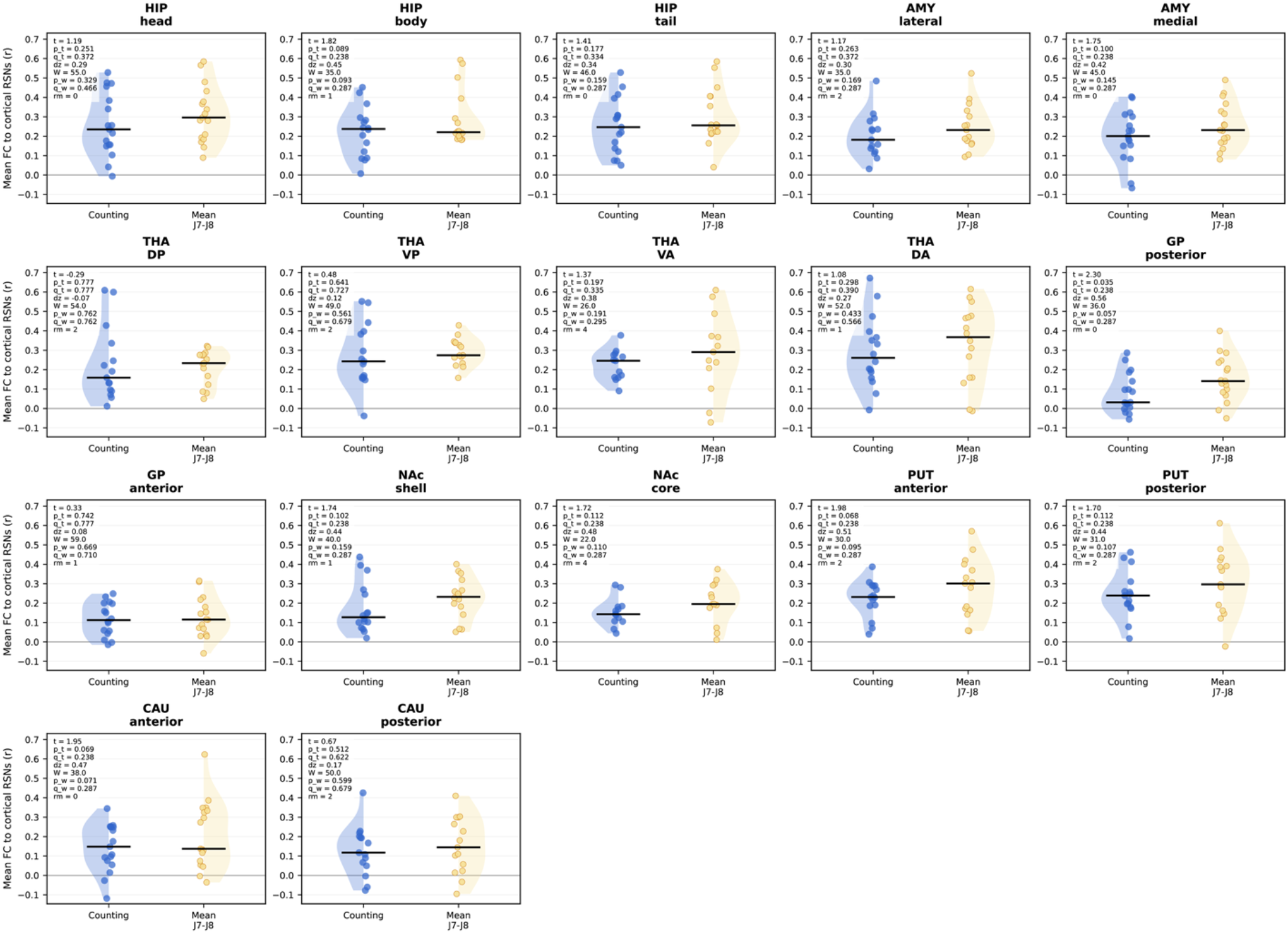
Subcortical-region connectivity to cortical resting-state networks during Counting and late Jhana. For each bilateral subcortical region, functional connectivity was first computed separately to each of the seven cortical RSNs and then averaged with equal RSN weighting, such that each cortical network contributed one value regardless of its number of nodes. Late Jhana was summarized as the within-subject mean of J7 and J8. Points show individual subjects, split violins show condition distributions, and black bars indicate medians. Statistics compare Counting vs mean J7-J8 after region-wise Tukey outlier trimming; annotations report paired t-tests, FDR-adjusted q-values across the 17 subcortical regions, Cohen’s d, Wilcoxon signed-rank tests, and the number of removed subject pairs. Most subcortical regions showed positive late-Jhana shifts, with the strongest raw increase in posterior globus pallidus, although no individual region survived FDR correction.

